# Increased layer 5 Martinotti cell excitation reduces pyramidal cell population plasticity and improves learned motor execution

**DOI:** 10.1101/2024.03.10.584276

**Authors:** Thawann Malfatti, Anna Velica, Jéssica Winne, Barbara Ciralli, Katharina Henriksson, George Nascimento, Richardson Leao, Klas Kullander

**Affiliations:** Department of Immunology, genetics and pathology, Uppsala University, Uppsala, Sweden; Brain Institute, Federal University of Rio Grande do Norte, Natal/RN, Brazil

**Keywords:** Martinotti alpha 2 cell, Assemblies, Motor function, Plasticity

## Abstract

During motor activity and motor learning, pyramidal cells in the motor cortex receive inputs from local interneurons as well as deeper structures. Layer 5 pyramidal cells in the primary motor cortex then feed commands to spinal circuits for motor execution. The genetic ablation of layer 5 Chrna2 Martinotti cells, which selectively target pyramidal tract pyramidal cells, resulted in disturbed fine motor functions. Using calcium imaging combined with chemogenetics, we show that activation of layer 5 Chrna2 Martinotti cells during training increases pyramidal cell tuning, changes responses temporal patterns and decreases assembly reconfiguration, while not affecting motor learning success rates. However, in mice that had already learned a reach-and-grasp (prehension) task, Chrna2 Martinotti cell activation resulted in improved prehension and increased power in low theta and high gamma bands of local field potentials in the motor cortex. This work indicates that activation of Chrna2 Martinotti cells reduces pyramidal cell assembly plasticity during learning, possibly facilitating already acquired motor skills.

## Introduction

Refined control of limb movement is crucial to perform essential behaviors such as building shelter, reaching for objects and exploring food. To initiate a voluntary movement, information is relayed from processed sensory information and command centers to circuits in the motor cortex, where the intended motor sequence is processed. The primary motor cortex (M1), which processes motor plans involving proximal and distal joints [1], delivers commands to spinal cord neurons to execute motor behaviors [2]. Motor plasticity continuously shapes and refines the neural circuits underlying motor control, ultimately influencing the execution of motor tasks in diverse contexts and conditions [3]. How the cortex is involved in motor learning and the execution of learned movements is still under debate. Most studies agree that cortical processing is necessary for the initiation of a learned motor sequence [4, 5, 6], but it remains unclear whether cortical processing is necessary for its execution. For example, a study used lesions and a lever pressing task to show that the rodent motor cortex is required for learning, but not for the execution of movements [7]. However, the use of optogenetic mid-movement activation of local inhibitory interneurons in the motor cortex could pause the execution of a learned movement, providing contradictory data [8]. Cortical excitatory and inhibitory interneurons serve as an interface between sensory inputs and motor commands and can thus provide entry points for the congregation of direct sensory input and processed information from the prefrontal cortex, to achieve the most purposeful movement at any given moment. Identifying the function and connectivity of specific interneuron populations within M1 is an important step towards disentangling the neuronal circuits that control and modulate motor function.

Martinotti cells (MCs) are inhibitory interneurons, mainly found in neocortical layer 3 and 5 [9], typically express the neuropeptide somatostatin (SST), and project to layer 1 where they form synapses onto the distal dendrites of cortical pyramidal cells [PCs; 10]. There are several studies investigating the connectivity and functional role of SST-expressing interneurons [11, 12, 13, 14, 15]. However, in the cortex, SST is expressed in a variety of diverse interneuron populations including large basket cells, double-bouquet cells, long-range GABAergic projection cells, bitufted cells, fanning-out MCs and T-shaped MCs [9, 16, 17]. In fact, only around 50% of SST interneurons are MCs [18] and MCs are found in all cortical layers except for layer 1 [19]. Recently it has been shown that more specific genetic markers can subdivide SST interneurons and MCs into subtypes that reside in different cortical layers and exhibit different morphologies and axonal projection patterns [20, 21], resulting in different layer specific functional properties [22, 16]. Indeed, a motor learning study has suggested that each cortical layer operates under unique constraints reflective of its specialized function [23].

In mice, a subset of MCs (herein referred to as M*α*2 cells) located in layer 5b, expressing the cholinergic receptor nicotinic alpha 2 subunit (Chrna2), send inhibitory projections selectively to thick-tufted type A pyramidal tract PC where they can reset and synchronize PC activity in a frequency-dependent manner [24, 21]. These findings suggest that M*α*2 cells are important for coordinated pyramidal tract PC activity, which is involved in top-down control of subcortical structures. However, how M*α*2 cells might influence PC activity to learn and execute purposeful motor behaviors is not understood. To probe the possible role of M*α*2 cells in motor processing, we evaluated skilled forelimb motor learning and execution in a prehension task during chemogenetic activation of M*α*2 cells, while simultaneously recording layer 5 PC activity using calcium imaging. We found that increased excitation of M*α*2 cells reduced plasticity in PC assemblies during performance refining, although learning efficiency was unaffected. Here, “assemblies” refer to functionally defined groups of pyramidal cells with temporally correlated activity patterns and are thought to represent cooperative network units underlying information processing. In contrast, motor performance of an already acquired skill was improved, suggesting that layer 5 Martinotti cells are required for cortical plasticity and motor execution.

## Materials and methods

### Animals

All animal procedures were approved by the local animal research ethical committee (Uppsala djurförsöksetiska nämnd) and followed the Swedish Animal Welfare Act (Svensk författningssamling (SFS) 2018:1192), The Swedish Animal Welfare Ordinance (SFS 2019:66) and the Regulations and general advice for laboratory animals (SJVFS 2019:9, Saknr L 150). Ethical permit number: 5.8.18-08463/2018, 5.8.18-08464/2018 and 5.8.18-07526/2023. Chrna2-Cre mice, produced and bred in our own facility [25], were crossed with either C57BL/6J (Taconic, Denmark) or GT(ROSA)26Sortm14(CAG-tdTomato)Hze [RRID:IMSR JAX:007914; 26] mice and the offspring were genotyped in house for the presence of the Chrna2-Cre allele and/or the tdTomato allele. The following primers were used: Chrna2-Cre 5’-gacagccattttctcgcttc-3’ (forward) and 5’-aggcaaattttggtgtacgg-3’ (reverse); tdTomato 5’-ctgttcctgtacggcatgg-3’ (forward) and 5’-ggcattaaagcagcgtatcc-3’ (reverse). The Chrna2-Cre, and tdTomato allele were kept heterozygous. Mice were housed with littermates in approximately 501 cm2 individually ventilated cages (up to 5 mice/cage) with bedding and enrichment (a carton house and paper tissues), kept in a 12-h light on/light off cycle (6 a.m.–6 p.m.), and maintained at 21 ± 2 °C with a humidity of 45-64%. Mice were provided food (diet pellets, Scanbur, Sweden) and tap water ad libitum, except for when food was restricted for behavioral testing (see single pellet prehension task and pasta handling task further down). After lens or electrode implant surgery, mice were housed individually to decrease risk of implant detachment. Mice were 6-25 weeks old at the start of experiments. Chrna2-Cre^tg/wt^ and Chrna2-Cre^wt/wt^ (Cre-negative littermates designated “controls”) were used. Chrna2-Cre^tg/wt^-tdTomato^lx/wt^ mice (and Cre-negative littermates, Chrna2-Cre^wt/wt^-tdTomato^lx/wt^; controls) were only used for experiments with Cre-dependent caspase3-induced apoptosis. Female mice were used in the behavioral experiments, except for in the miniscope recordings in which male mice were used. Throughout the paper, control mice refer to Cre-negative littermates that have received the same treatment as Cre-positive mice.

### Viral injections

Mice were sedated with 1-4% Isoflurane (Baxter). Analgesia was given subcutaneously; 2 mg/kg bupivacaine (Marcain, AstraZeneca), 5 mg/kg carprofen (Norocarp vet, N-vet or Rimadyl Biovis vet) and 0.1 mg/kg buprenorphine (Vetergesic vet, Ceva). A midline incision was made in the scalp, muscles and periosteum were removed using 3% hydrogen peroxide (Sigma-Aldrich) and a hole (1 mm in diameter) was drilled at the site of injection (coordinates relative to bregma: AP 0.8 mm, ML 1.5 mm and DV −1.3 mm) with a hand-held drill. The injection coordinates targeted the forelimb area of the M1. The injection coordinates were chosen based on the Mouse Brain Atlas [27], as well as previous studies mapping the motor cortex [28, 29]. Viral vectors (for more information see below for each experiment) were injected using a 10 µl Nanofil Hamilton syringe (WPI, USA) mounted on a stereotaxic frame. Ten minutes post injection the needle was slowly withdrawn from the brain and the wound was stitched with resorbable sutures (Vicryl rapide, Ethicon, 6-0). Mice were then left to recover for 2-4 weeks before any behavioral experiments were initiated. Viral transduction was confirmed by post-hoc histological analysis of brain sections.

For Cre-dependent caspase3-induced apoptosis, 14 female mice (7 Chrna2-Cre^tg/wt^-tdTomato^lx/wt^; 7 Chrna2-Cre^wt/wt^-tdTomato^lx/wt^) were injected bilaterally with 750 nl of AAV5-FLEX-TACASP3-TEVP (Addgene viral prep # 45580-AAV5, viral titer: 4.6 x 1012 VG/ml) at a speed of 100 nl/min and were used for behavioral experiments. Two additional mice were injected unilaterally in order to check the percentage of neuronal ablation comparing the injected to the non-injected side.

For chemogenetic activation during local field potential (LFP) recordings, 14 female mice (7 Chrna2-Cre^tg/wt^; 7 Chrna2-Cre^wt/wt^) were injected bilaterally with 300 nl of AAV9-hSyn-DIO-hM3Dq-mCherry (Addgene viral prep # 44361-AAV9, viral titer 2.3 x 1013 GC/ml) diluted in 300 nl sterile saline (9 mg/ml, Fresenius Kabi, Sweden) at a speed of 200 nl/min.

For repetitive chemogenetic activation during training of the single pellet prehension task, 20 mice (10 Chrna2-Cre^tg/wt^; 10 Chrna2-Cre^wt/wt^) were injected bilaterally with 300 nl of AAV9-hSyn-DIO-hM3Dq-mCherry diluted in 300 nl sterile saline (9 mg/ml, Fresenius Kabi, Sweden) at a speed of 200 nl/min.

For calcium imaging of pyramidal cell activity, 8 mice (5 Chrna2-Cre^tg/wt^; 3 Chrna2-Cre^wt/wt^) were first injected bilaterally with 300 nl of AAV9-hSyn-DIO-hM3Dq-mCherry diluted in 300 nl sterile saline (9 mg/ml, Fresenius Kabi, Sweden) at a speed of 200 nl/min. One week post injection, the mice were injected unilaterally in the right hemisphere with 600 nl of AAV9-CamKII-GCaMP6f-WPRE-SV40 (Addgene viral prep # 100834-AAV9), at a speed of 200 nl/min. One week post-injection of AAV9-CamKII-GCaMP6f-WPRE-SV40, the mice underwent surgery for lens implantation (see Lens implantation below).

### Electrode array assembly and implantation

For LFP experiments, mice were injected with AAV9-hSyn-DIO-hM3Dq-mCherry (see viral injections above) bilaterally three weeks before electrode implantation surgery. Tungsten insulated wires of 35 *µ* m diameter (impedance 100-400 k Ω, California Wires Company) were used to manufacture 2×5 arrays of 10 tungsten wire electrodes. The wires were assembled to a 16 channel custom made printed circuit board and fitted with an Omnetics connector (NPD-18-VV-GS). Electrode wires were spaced 200 *µ* m and the array was implanted into the right hemisphere of the M1. Surgery preparation was performed as described for viral injection, and four small craniotomies were done in a square at coordinates AP 0.70 mm, ML 1.25 mm; AP 0.70 mm, ML 2.25 mm; AP 1.0 mm, ML 1.25 mm; and AP 1.0 mm, ML 2.25 mm; to make a cranial window were the electrodes were slowly inserted at DV −1.30 mm. Four additional craniotomies were drilled for the placement of anchoring screws (AgnThos, MCS1×2), where the screw placed over the cerebellum served as reference. The electrode implant was fixed to the skull with dental cement (Tetric EvoFlow) around the anchor screws. After surgery, the mice were monitored until awake, housed individually and allowed to recover for one week before recordings.

### Lens implantation

The procedure was initially identical to the viral injections, but instead of injecting a virus, a prism lens (CLH type glass, lens diameter 1.0 mm, prism size 1.0 mm, total length of lens+prism 3.68 ± 0.156 mm, working distance 0.3 mm, enhanced aluminum and SiO2 protective coating, Go!Foton, USA) combined with a relay GRIN lens (Imaging Focusing Rod Lens, diameter 2.0 mm, working distance at infinity) for recordings of cortical layer 5 was implanted unilaterally using the stereotaxic frame. To implant the prism lenses, the tip of a dissection knife was inserted 0.3 mm into the cortex for three minutes before the lens was lowered into the cortex with the tip 0.3 mm lateral and −0.5 mm dorsoventral to the viral injection coordinates. The lens was fixed to the skull and anchor screws with cyanoacrylate glue (OptiBond, Kerr Dental). Silicon glue (Kwik-Sil, World Precision Instruments) was added to protect the lens from mechanical damage in between experiments. Mice were treated with Enrofloxacin 5 mg/kg (Baytril) for two days pre-surgery and seven days post-surgery. Two to three weeks post implantation the mice were again anesthetized, the scalp incised, and the baseplate (used to attach the miniscope during recordings) was mounted on top of the skull with cyanoacrylate glue. To verify that the prism lens was correctly positioned, we performed post-hoc histological analysis to confirm that it was centered around the cortical layer 5.

### Drug preparation

The most commonly used chemogenetic agonist is clozapine-N-oxide (CNO), however, it has been suggested that the main effect on the engineered receptors, after systemically administered CNO, is mediated by back-converted clozapine [CLZ; 30, 31]. Since the conversion rate might differ between individuals [32], we chose to use low dose CLZ for chemogenetic experiments. A stock solution was made from 1 mg of CLZ (Hello Bio, batch E0697-1-1) dissolved in 40 µl dimethyl sulfoxide (DMSO, Sigma-Aldrich). The stock solution was kept at room temperature for the entire set of experiments. The working solution given to the mice was made fresh daily; 2 µl of the stock solution were diluted in sterile saline to a final drug concentration of 1 µg/ml. The mice then received 0.01 mg/kg CLZ intraperitoneally (i.p.).

### Single pellet prehension task

The single pellet prehension protocol consisted of handling, habituation and test sessions. The handling and habituation were performed the same way throughout all experiments, while the number of test sessions varied based on the experimental design (see below for more information about each experimental group). During handling, each mouse was placed on the body and hands of the experimenter for five minutes during two consecutive days and sugar pellets (non-pareille, Dr.Oetker) were inserted in the home cages after handling. For habituation, after completed handling, mice were placed inside the test arena for ten minutes and allowed to freely behave with sugar pellets on the floor of the arena during two consecutive days. The arena consisted of an acrylic rectangular box of dimensions 22 cm x 9 cm x 20.5 cm with a vertical slit (0.5 cm wide) in one of the narrower walls. An elevated (2cm) plate was attached to the outer wall of the arena, 2cm to the right of the slit. On the third day of habituation, mice were presented sugar pellets on a spoon, outside the slit of the arena, which they were allowed to retrieve with their tongues. Following habituation, the mice underwent ten minutes long test sessions every other day. All test sessions were performed after 15-20 hours of food restriction, starting at 5 p.m. the previous day. After the test session, the mice were returned to their homecage with free access to food and water. For all mice, the initial sessions were performed with spoon aided pellet delivery, to make it easier for mice to understand that they should reach for the pellet. The spoon was initially placed closer to the opening of the slit to increase success rate and motivation to perform the test. As the mice reached more, the spoon was gradually moved further away from the slit untill it was at the same distance as a fixed position on the plate outside the slit. After the fifth session, the pellet was presented on the fixed position on the plate, instead of on the spoon. The use of a gradually increasing difficulty in the task is needed due to the very challenging position of the plate. The plate was placed 2cm to the right of the slit opening, forcing the mice to reach with their left paw. The female experimenter was blinded regarding mouse genotype during the test. The behavior was recorded from the front using a high-speed camera (Panasonic high-speed camera, HC-V750) at 1920×1080 px fixed at 50 FPS.

To clarify the session structure: sessions 1–5 were performed with pellet delivery using a spoon, whereas from session 6 onward pellets were presented on a plate positioned at the slot. During sessions 1–5, pellet placement was progressively adjusted to increase task difficulty (i.e., gradually positioned closer to the plate slot), thereby shaping forelimb reaching behavior. In session 6, the pellet was placed directly on the plate slot, requiring the animal to adapt its previously acquired reaching strategy. For the caspase3 induced apoptosis group, after completing handling and habituation, 10 mice (4 Chrna2-Cre^tg/wt^-tdTomato^lx/wt^ and 6 Chrna2-Cre^wt/wt^-tdTomato^lx/wt^) were trained to take pellets from a spoon through the slit during four sessions, then trained to take pellets from the fixed position on the plate during three sessions. One mouse did not reach for pellets on the seventh session, and was therefore excluded. 9 mice (3 Chrna2-Cre^tg/wt^-tdTomato^lx/wt^ and 6 Chrna2-Cre^wt/wt^-tdTomato^lx/wt^) were selected and received viral vector injections (AAV5-FLEX-TACASP3-TEVP), followed by five more test sessions, starting at four weeks post-injection.

For the LFP group, 11 mice (5 Chrna2-Cre^tg/wt^; 6 Chrna2-Cre^wt/wt^) were used. After completing handling and habituation, the mice were trained during four sessions (session 1-4) to take pellets from a spoon through the slit, then trained for three sessions (session 5-7) to take pellets from the fixed plate through the slit. One mouse that did not reach for pellets in the seventh session was excluded. 13 mice (6 Chrna2-Cre^tg/wt^ and 7 Chrna2-Cre^wt/wt^) were selected and received viral vector injections (AAV9-hSyn-DIO-hM3Dq-mCherry), followed by six test sessions (session 8-13) with the headstage connected for LFP recordings four weeks post injections The LFP signal was recorded with a 16-channel Intan RHD 2132 headstage connected to an Intan RHD2132 amplifier board by a thin flexible wire. Mice were briefly anesthetized with isoflurane in order to connect the headstage, then allowed to fully recover and placed inside the arena. Afterwards, mice were disconnected from the headstage and returned to their homecage. In the first three sessions after viral injections (session 8-10) the mice received 0.9 % NaCl i.p. 45 minutes before the start of the recording. In the last three sessions (session 11-13) the mice received 0.01 mg/kg CLZ i.p. 45 minutes before the start of the recording.

For the group with repetitive M*α*2 cell activation during learning, 20 mice (10 Chrna2-Cre^tg/wt^; 10 Chrna2-Cre^wt/wt^) were injected with viral vectors (AAV9-hSynDIO-hM3Dq-mCherry). Three to four weeks after viral injection, the handling and habituation was started. 45 minutes before the start of each session, the mice were injected with 0.01 mg/kg CLZ i.p. The first five sessions, the mice reached through the slit for pellets placed on a spoon. After five minutes in the fifth session, the pellet was instead placed on a fixed position on the plate, and the mice were encouraged to reach for it for five more minutes. The following five sessions, the mice reached for the pellet in the fixed position on the plate during ten minute long sessions. Mice with no reaches in more than three sessions were excluded from the analysis, which resulted in exclusion of one Chrna2-Cre^tg/wt^ mouse and three Chrna2-Cre^wt/wt^ mice. Three Chrna2-Cre^tg/wt^ mice were excluded due to no DREADD expression on post hoc tissue analysis.

For the miniscope group, 8 mice (5 Chrna2-Cre^tg/wt^; 3 Chrna2-Cre^wt/wt^) underwent viral injections and lens implant. 2 Chrna2-Cre^tg/wt^ did not have any detectable expression of the calcium indicator GFP Calmodulin M13 peptide 6 fast (GCaMP6f; checked with the miniscope under anesthesia before baseplate fixation), so they were excluded before the baseplate fixation step. The handling was started two to five weeks after lens implantation, followed by one additional habituation step in which the mice were habituated to the miniscope. The mice were then sedated with 1-4% isoflurane for two to five minutes while the miniscope was mounted on the baseplate and thereafter placed in a circular arena (diameter 45 cm) for 45 minutes, then returned to their homecage for 45 minutes and then once again placed in the circular arena for 45 minutes. After this habituation step the mice were sedated with 1-4% isoflurane for two to five minutes while the miniscope was disconnected from the baseplate. The mice were then returned to their homecage. A few days later, the mice underwent habituation to the single pellet prehension task as previously described, followed by six test sessions. 45-60 minutes before each test session the mice received 0.01 mg/kg CLZ i.p.. The miniscope was mounted on the baseplate on session 1, 2, 5 and 6. Only on those sessions, approximately 20 minutes after CLZ administration, the mice were sedated with 1-4% isoflurane for two to five minutes while the miniscope was mounted on the baseplate. After 20 minutes recovery from the sedation, the mice were placed in the prehension arena for ten minutes. In the arena, the cellular activity was recorded with the miniscope on sessions 1, 2, 5 (pellet delivered on the spoon) and 6 (pellet placed on the fixed plate) and the behavioral activity was recorded with the video camera on session 1-6. A blinking LED was recorded with the behaviour video camera to enable synchronization of the time stamps of the miniscope recordings with the behavioral videos. After the test, the mice were sedated with isoflurane in order to remove the miniscope from the baseplate. Session 1-5 were performed every other day, session 6 was performed on the same day as session 5 directly after session 5. Data was recorded using a data acquisition device (DAQ) and a computer with Miniscope controller software installed. Focus of the miniscope was adjusted manually (300 *µ* m range). Optimal laser power (around 30% of max brightness), imaging gain, and focal distance was selected for each mouse and conserved across all sessions. The calcium signal was acquired at a frame rate of ≈ 20 frames per seconds, with the timestamp of each frame registered onto a separate file.

### Pasta Handling Test

The mice were tested in the pasta handling task, a quantitative assessment of skilled forelimb function, in order to measure forepaw dexterity [adapted from 29]. Mice were exposed to the pasta pieces (uncooked capellini pasta, 1.0 mm diameter, 2.6 cm length) in their home cage for five days to avoid neophobic reactions. Thereafter, the mice were habituated to eat inside the test arena (22 cm x 9 cm x 20.5 cm) for three days, without food restriction. After the habituation period, the mice were tested in the pasta handling test. The test consisted of sequential trials in which one pasta piece was delivered on the floor, in the center of the arena, for each trial. The behavior was recorded from the front using a high-speed camera (Panasonic high-speed camera, HC-V750) at 1920×1080 px fixed at 50 FPS. Test sessions were performed after 15-20 hours of food restriction, starting at 5 p.m. the previous day. Each session of ten minutes consisted of a maximum of five trials. To make it easier to see how the capellini strand would move while being consumed, each pasta piece received a food coloring mark on one half. The number of pasta drops and the time spent handling the pasta was then quantified by visual inspection. 9 mice (3 Chrna2-Cre^tg/wt^-tdTomato^lx/wt^; 6 Chrna2-Cre^wt/wt^-tdTomato^lx/wt^) bilaterally injected with AAV5-FLEX-TACASP3-TEVP were tested in the pasta handling task six weeks after viral injection and one week after the last pellet prehension task session.

### Hanging Wire Test

The coordination and strength of the forepaws was tested by the hanging wire test, in which mice suspend their body by holding on to a single wire stretched between two posts 40 cm above the ground. The mice were placed on the middle of the wire hanging from its forepaws for three minutes. A paper bed was placed between the two posts to avoid injury when the mice fell. The time hanging on the wire and number of falls were quantified by visual inspection. 9 mice (3 Chrna2-Cre^tg/wt^-tdTomato^lx/wt^; 6 Chrna2-Cre^wt/wt^-tdTomato^lx/wt^) were used. Mice were injected with viral vectors (AAV5-FLEX-TACASP3-TEVP) bilaterally, then performed the single pellet prehension task, and after completion of the single pellet prehension task, they performed the pasta handling and finally the hanging wire test (six weeks after the viral vector injection).

### hM3Dq activation validation

Six weeks after the injection of AAV9-hSyn-DIO-hM3Dq-mCherry (n = 9 Chrna2-Cre+ and 5 controls), CLZ 0.01 mg/kg was administered i.p. 135 minutes before transcardial perfusion and tissue preparation. After the administration of CLZ, before the transcardial perfusion, the mice were kept in their homecage. The mice were then deeply anesthetized with Medetomidin 1mg/kg (Dormitor Vet, Orion Pharma Animal Health) and Ketamin (Ketalar, Pfizer) 75 mg/kg i.p., before they were transcardially perfused as described below. After dissection and sectioning, immunohistochemistry for c-Fos and mCherry was performed as described below.

### Tissue preparation and immunohistochemistry

To fix the tissue for post-hoc analysis, mice were anesthetized with Medetomidin 1mg/kg (Dormitor Vet, Orion Pharma Animal Health) and Ketamin (Ketalar, Pfizer) 75 mg/kg i.p. and sacrificed by transcardial perfusion with phosphate buffered saline (PBS, Fisher BioReagents, CAT 10051163) followed by 4% formaldehyde (VWR Chemicals BDH^®^, CAT 9713.1000). The brains were dissected out and post-fixed in 4% formaldehyde overnight at 4°C, washed three times for ten minutes in PBS and then kept in PBS at 4°C until sectioning (a few hours or days). Before they were sectioned, the brains were embedded in 4% agarose in PBS. Sections were cut at a thickness of 70 µm using a Vibratome (Leica VT1000S) and then mounted with ProLongTM Gold Antifade Mountant (ThermoFisher Scientific, CAT P10144) or Fluoroshield (Abcam) on glass slides (Superfrost^®^Plus, Thermo Scientific).

For the mice used in the hM3Dq activation validation, immunohistochemistry was performed. After sectioning these brains, the sections were washed in PBS and pre-blocked for one hour in 2% donkey serum (Biowest), 1% Bovine Serum Albumin (BSA, Sigma Aldrich), 0.1 % Triton^®^X-100 (Sigma Aldrich) and 0.05% Tween^®^20 (Sigma Aldrich) in PBS. The primary antibody, c-Fos(4)-G (goat polyclonal IgG, Santa Cruz Biotechnology, CAT K1314) or c-Fos (9F6) (Cell signaling, rabbit, #2250) and anti-mCherry (ThermoFisherScientific, 16D7, rat, M11217) were diluted 1:500 in 0.25% gelatin and 0.5 % Triton^®^X-100 in Tris-HCl buffered saline (TBS; 0.3% Tris, 0.8% NaCl and 0.02% KCl in mqH2O, pH 7.4) and added to the sections. After 48 hours of incubation at 4°C the sections were washed with TBS and the secondary antibodies, donkey anti-goat A647 (ThermoFisherScientific A21447), donkey anti-rabbit A647 (ThermoFisherScientific A31573) and donkey anti-rat A488 (ThermoFisher-Scientific A21208) were diluted 1:500 in 0.25% gelatin and 0.5 % Triton^®^X-100 in TBS, added to the sections and incubated for 90 minutes at room temperature. The sections were washed with 0.1% Tween in TBS and distilled water before they were mounted on glass slides.

### Imaging and image post processing

Most wide field images were acquired on a Zeiss Axio Imager Z2 (Zeiss, Germany) with a 10× objective (effective NA 0.45), a colibri LED 7 and Zen blue software (Zeiss, Germany), using a multi-band bandpass filter (Zeiss, Germany) with the filter excitation wavelengths 370-400 nm, 450-488 nm, 540-570 nm, 614-647 nm, 720-750 nm and the filter emission wavelengths 412-438 nm, 501-527 nm, 582-601 nm, 662-700 nm, 770-800 nm for the fluorophores DAPI, Alexa 647 and GFP and a single-band bandpass filter (TxRed-4040C-ZHE-ZERO, Semrock) with the excitation wavelengths 540-552 nm and emission wavelengths 590-4095 nm for mCherry. The light source used for the different fluorophores was LED-module 385nm for DAPI, LED-module 630 nm for Alexa 647, LED-module 475 nm for GFP and LED-module 567 nm for mCherry. Some images were taken using an Olympus BX61WI fluorescence microscope (Olympus, Japan) with a 10× (effective NA 0.40) objective and a Volocity 4.1.0 software (Quorum Technologies) or a Zeiss LSM700 confocal microscope (Bio-Vis facility, Uppsala University). Brightness and contrast were adjusted in the Fiji software [33], equally for the whole image and without obscuring any data.

### Pellet prehension task analysis

For each pellet prehension task session, the positions of the pellet, both forepaws, thumb, index and middle fingers were tracked. The AI network used for tracking (see Software availability) was trained specifically for detection of the left paw and the three fingers at the prehension onset position (paw palm pointing inside, thumb pointing up and fingers pointing to the medial line), and a tracking likelihood threshold of 70% was achieved from prehension onset to grasp end. Therefore, prehensions were reliably detected as blocks where the tracking likelihood for the left paw and its three fingers was above 70%. Prehensions were classified as successful when the animal reached for the pellet, grasped the pellet and brought it to its mouth. Any other outcome (mice hitting the pellet but failing to grasp it; missing the pellet; grasping the pellet but dropping it before it reached the mouth; reaching for the pellet when no pellet was available) was considered a failed prehension.

### LFP electrophysiology analysis

Square voltage pulses were used to trigger LFP and miniscope recordings; and to drive a red LED placed at the top right corner of the behavioral camera field of view. Therefore, behavioral video recordings were synchronized with the corresponding electrophysiological recording by detecting the LED onset in the video. LFP data for each channel was filtered using a second-order Butterworth bandpass filter across the entire frequency range of 1-100 Hz, as well as within specific bands: theta (4-12 Hz), gamma (30-90 Hz), low theta (4-7 Hz), high theta (8-12 Hz), low gamma (30-40 Hz), and high gamma (60-90 Hz). This process was followed by the computation of the spectrogram, which offers a time-resolved representation of the frequency content in the data. The resulting spectrogram was normalized by dividing each power value by the cumulative mean, ensuring that the data reflected relative power rather than absolute magnitudes. Subsequently, average power values were extracted within the designated frequency bands by averaging over the frequency dimension, focusing specifically on the frequency bins that fell within the defined limits of each band.

### In-vivo calcium imaging analysis

#### Neural signal extraction

The calcium signal in PCs was obtained using the viral vector with CamKII promoter driven expression of GCaMP6f. The increase in calcium levels corresponds to increased cell activity, and single spikes can only be indirectly estimated [34, 35, 36]. Miniscope videos were spatially downsampled 3x and motion-corrected, then cells signal was extracted using constrained nonnegative matrix factorization for microendoscopic data (CNMF-E) approach [37], and fluorescence traces were detrended by baseline subtraction. The baseline components were estimated using a running 8th percentile derived from kernel density estimation over a 250-frame window, and subtracted from each respective trace. Since true F0 estimation is not possible on 1-photon data, the baseline-subtracted trace is here referred to as ΔF detr.. Neuron identity was matched along multiple recordings through intersection over union metric and the Hungarian algorithm for optimal matching. Up to this step, all miniscope video analysis was done as implemented previously [38].

#### Neural activity around prehensions

Prehension onset (time = 0 ms in all plots) was defined as the moment when the mouse’s paw forms the grasp position, flipping the palm inside, thumb pointing up and fingers pointing to the medial line, then neurons fluorescence traces were sliced 1 s before and 1 s after forepaw prehension onset, and slices were grouped by prehension accuracy (successful or failed) and averaged separately for each session. To specifically assess how M*α*2 cell activity modulates pyramidal cell (PC) dynamics during prehension, we restricted our analyses to PCs that showed significant task-related modulation. For each neuron, we verified whether calcium activity is different between the before onset and after onset phases (ANOVA or Kruskal-Wallis, according to normality of distribution, evaluated using Shapiro’s test). Only neurons that displayed a significant change in activity between these epochs were included in subsequent population, temporal, and assembly analyses. Neural activity was evaluated independently for successful and failed prehensions, and neurons with a significant change in at least one of the accuracies were included for each session. This approach ensured that our dataset was enriched for neurons that participate in prehension-related processing, rather than including large numbers of non-task-related cells that would dilute movement-specific effects. Using this criterion, 15.97% of recorded PCs were excluded because they did not show significant prehension-related modulation.

Pyramidal cell amplitude was calculated as the mean fluorescence of the averaged prehension window slices. For L5 pyramidal cells fluorescence peak latency analysis, traces for successful or failed grasps were separately averaged and normalized for each neuron, and neurons were sorted according to peak latency at successful grasps from each session. For PC response peak width, the width in seconds of the maximum response peak was calculated at half the peak prominence.

#### Assembly analysis

Additional constraints were required to allow for detection of assemblies between sessions. This limited the analysis to the sessions 1, 2 and 6, and only PCs detected through those sessions whose identity could be matched along sessions were kept. Therefore, on average 84.8 ± 14.2 PCs (n = 4 mice; 2 ctrl, 133 and 68; 2 Chrna2-Cre+, 77 and 61) were kept. Assemblies were identified during the entire 10 minute recording of a session, such that assembly detection was not affected by prehension accuracy. Assemblies were detected using independent component analysis with circular shifts [ICA-CS; 39], as implemented previously [40]. After removing neurons with no significant change comparing the before onset and after onset phases, only assemblies with 4 or more neurons and only neurons belonging to at least one assembly were kept for further assembly analysis. For calculating assembly spatial distribution, neurons from the assembly were first sorted according to their cartesian angle relative to the center of the assembly, which is the average of neurons’ (x,y) coordinates. Finally, the assembly distribution was calculated as the area of the polygon formed by the sorted neurons. Assemblies resilience was calculated by finding the assembly with highest neuronal composition intersection on the other sessions according to the following formula:

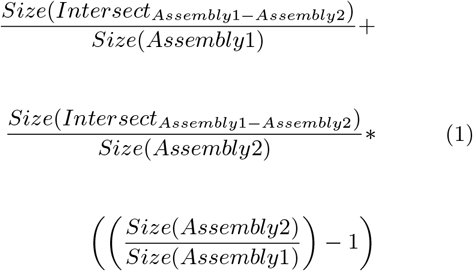

such that resilience is within 0 (no intersection between assemblies) and 1 (exactly the same neurons forming both assemblies). To find assembly activity peaks, 1000 shuffles of activity of all neurons within an assembly on the time dimension were done, and activity peaks were considered as time samples where the sum of all neurons’ real activity is larger than 95% of the sum of the shuffled neuronal activity, resulting in a rasterized activity (containing only 0 or 1 meaning inactive or active for each time point). Event histograms were calculated by slicing the assembly activity peaks in time windows of 2 s around each prehension (1 s before and 1 s after onset), where each bin corresponds to a one time sample of 50 ms, limited by the miniscope’s 20 FPS recording speed. Assembly salience, a measurement of how relevant the activation (or silencing) of an assembly is within the prehension movement, was calculated as

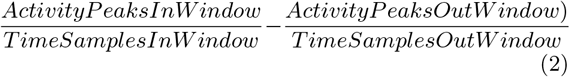

such that salience is within −100 (assembly is active every time sample out of the prehension window and not active on any time sample within the window, i.e., absent prehension-related activity and full activity outside the prehension window) to 100 (assembly active in all time samples within the prehension window and at no time sample outside the window, i.e., full prehension-related activity and absent activity outside the prehension window).

#### Trial adjacency interference

To address whether closely spaced prehension attempts influenced baseline neural signals, we evaluated whether trials with inter-trial intervals ≤ 2 s (the full prehension window duration) showed significantly different calcium activity between “previous” and “current” trials. We determined if neural responses were significantly modulated by prior trials within this window using ANOVA or ANOVA-type statistics [41], based on normality testing via Shapiro’s test. False discovery rate (FDR) correction was applied to p-values derived across all trial pairs using the Benjamini-Yekutieli procedure.

To further evaluate whether trial interference affects baseline neural signals, we analyzed fluorescence traces from time windows (−5 to −3 s) and (−3 to −1 s) relative to prehension onset, including only trials not preceded by another trial for at least 6 s. Fluorescence traces were sliced 5 s prior to 1 s prior to prehension onset, and grouped by genotype, training session and prehension accuracy (successful or failed) and averaged separately. An effect of time window epoch ((−5, −3) vs (−3, −1)) was determined via ANOVA or ANOVA-type statistics tests based on normality testing with Shapiro’s test and used to identify baseline interference effects.

### Statistical analysis

Differences were evaluated using mixed-models ANOVA, with post-hoc t-test for pairwise comparisons. Where the data showed non-normal distribution, evaluated using Shapiro’s test, ANOVA-type statistics [41, mixed models] or Scheirer-Ray-Hare (independent factors) were used for multiple factors; Kruskal-Wallis with F-ratios [42] and ANOVA-type statistics tests were used for independent and dependent single factors, respectively; and post-hoc Mann–Whitney U or Wilcoxon signed-rank tests for independent and paired factors, respectively. All possible pairwise comparisons were computed, with multiple comparisons adjusted by Holm correction, and statistical significance conditioned to a false positive risk lower than *α*= 0.05. In all figures, significance bars indicate significant comparisons (p *<* 0.05). The absence of these bars for any comparison means no significant difference was found (p *>* 0.05). Significance bars without ticks represent pairwise comparisons, while significance bars with downward ticks represent an effect. Error bars represent standard error of the mean (s.e.m) for all figures. For all violin plots, the filled area represents the entire data range, horizontal lines and triangle markers represent medians and means, respectively.

### Software availability

LFP recordings were done using the Open-ephys GUI [43]. Calculations were done using Scipy [44], Numpy [45] and SciScripts [46], all plots were produced using Matplotlib [47] and schematics were drawn using Inkscape [48]. Cell fluorescence signals were extracted and processed from miniscope videos using CaImAn [38] and OpenCV [49]. Tracking of the mouse forepaw and fingers was done using DeepLabCut [50]. All scripts used for data analysis, including the DeepLabCut trained network, are available online [git repository; 51].

## Results

### Increased M*α*2 cell excitation affected calcium signal amplitudes in PCs during training and retraining

Considering the dual roles of the M1 as a motor control structure and a dynamic substrate that participates in motor learning [52], we were interested in how PCs were affected by modulation of M*α*2 cells during motor learning. For this purpose, we turned to chemogenetic manipulation using designer receptors exclusively activated by designer drugs [DREADDs; 53]. In pilot studies, we found that increased excitation of M*α*2 cells, but not increased inhibition, gave a measurable effect in a motor test (using modified Gq-coupled human M3 or Gi-coupled human M4 muscarinic receptors [hM3Dq/hM4Di; 54]. We thus chose to continue with hM3Dq, and we first validated the effect of clozapine (CLZ) in Chrna2-Cre+ mice. To validate the clozapine (CLZ) induced activation of hM3DGq+ cells, we injected viral vectors carrying the modified Gq-coupled human M3 muscarinic receptor (hDM3Gq) and the fluorescent marker mCherry [55] under Cre-dependent expression and performed immunolabeling for c-Fos, a proto-oncogene expressed in most neurons after depolarization [56, 57, Supplemental Figure 1A-B]. As expected, hDM3Gq activation induced c-Fos expression in Chrna2-Cre+ mice (n=9; number of sections=346), on average, 32 mCherry-positive cells per hemisection were found, of which 76% were c-Fos positive. In control mice (n=5, number of sections=60), on average 5 mCherry positive cells per section were found, of which 0.8% were c-Fos positive (Supplemental Figure 1C). Throughout the manuscript, control mice refers to Cre-negative littermates that received the same treatment as the experimental Chrna2-Cre+ mice. All calcium imaging recordings were performed with mice under clozapine 0.01mg/kg treatment.

Next, a prehension task, in which mice are encouraged to reach for and grasp a sugar pellet, was used to evaluate fine motor functions and motor skill refinement [58, 59, 60]. Most dendritic spines that correlate with learning are formed after the initial training session and are maintained in the second training session [61]. Thus, we performed a set of recordings during the first session for collection of data in untrained mice, the second session and fifth sessions to follow performance refinement, and finally in session 6, during which a more challenging presentation of the pellet was introduced (Figure 1A-C). The contribution of increased M*α*2 cell excitation on individual PCs was investigated with calcium imaging. We recorded GCaMP6f fluorescence changes in cortical layer 5 using an implanted optical lens and prism assembled to a miniscope [Figure 1A; 62, 63]. To maximize the number of cells included in the analysis, PCs were separately identified for each session. Therefore, the number of PCs per animal varies for each session. On average, 61.4 ± 11.49 PCs (n = 6 mice; 3 Control, total of 1070 cells for 4 sessions, average of 105.2, 102.8 and 59.5 for each animal; 3 Chrna2-Cre+, total of 403 cells for 4 sessions, average of 45, 53.8 and 16) were included in the analysis. Calcium transients from individual cell activities were readily identifiable and showed diversified activity profiles (Figure 1D-E, Supplemental Figure 2A-H). Our analysis revealed no effect of genotype on prehension performance in a separate group of mice treated with CLZ before each session (n = 13 mice; 6 Chrna2-Cre+ and 7 controls; Supplemental Table 1A-C; Figure 1F-H). However, mice display improvement in the task (overall effect of training session for total number of prehensions, number of successful prehensions, and success ratio, Supplemental Table 1D-F), with an increase in the average number of total prehensions and successful prehensions from the first to the second session, as well as an expected decrease in the fifth (harder spoon session) and sixth sessions (pellet on the plate) due to increased difficulty and stable performance after session 7. The number of successful attempts per minute is an alternative measurement of performance progression, as the absolute number of attempts can increase disproportionately compared with the number of successful attempts, temporarily decreasing the success ratio even though the overall motor performance is still improving [59].

**Figure 1.**
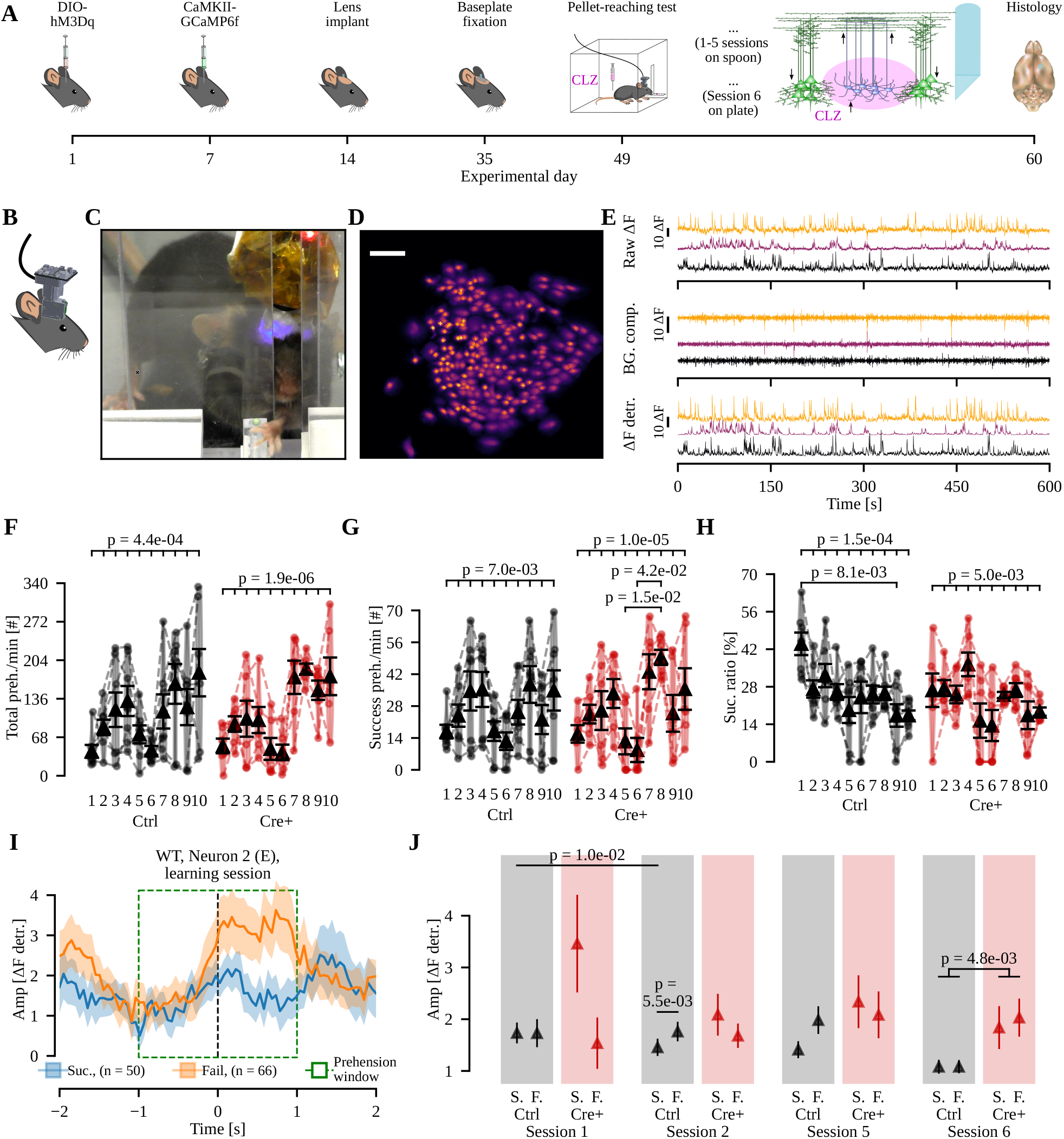
Mice with increased M*α*2 cell excitability displays increased PC activity after training and upon retraining. A) Timeline of experiments showing viral injections, miniscope lens implantation, base plate fixation and pellet-reaching task. GCaMP6f fluorescence and forepaw movements were recorded in sessions 1, 2, 5 and 6. B-C) Schematics (B) and photograph (C) of a mouse with the miniscope connected during prehension. D) Representative miniscope spatial footprints of L5 pyramidal cells after cell detection. Scalebar: 50 µm. E) Three representative example fluorescence traces from PCs showing the raw ΔF (top), estimated background components (middle) and the detrended traces (bottom). F-H) Total number (F), success number (G) and success ratio of prehensions (H), separated by session and genotype (Controls, ‘-’, black; Chrna2-Cre+, ‘+’, red). n mice = 7 Controls and 6 Chrna2-Cre+ (ANOVA). I) Representative traces from the second neuron from (E) showing the ΔF detr. activity −2 to 2 second around the prehension onset, averaging all successful (Suc., blue) and failed prehensions (Fail., orange). J) Average amplitude of PC activity represented as mean *±* SEM, separated by session (1, 2, 5, 6), genotype (Controls, ‘Ctrl’, black; Chrna2-Cre+, ‘Cre+’, red) and prehension accuracy (S., successful prehensions; F., failed prehensions). n cells per session = 162, 363, 399, 146 for Controls; 106, 126, 53, 118 for Chrna2-Cre+ (Scheirer-Ray-Hare followed by Mann-Whitney U).

As expected, increasing M*α*2 cell excitation led to a decrease in baseline PC activity (non-prehension related) in Chrna2-Cre+ mice compared to controls (Supplemental Figure 2I). When looking specifically at PC activity 2s around the prehension onset (−1 to 1s), our results showed an effect of session in PC amplitude on successful prehensions of controls (Supplemental Table 2A-C). PC activity amplitudes were decreased during successful prehensions in the second session compared both to failed prehensions (Supplemental Table 2D) and to successful prehensions from the first session (Supplemental Table 2E; Figure 1I-J). Furthermore, an overall effect of genotype on PC activity amplitude was found for session 6 (Supplemental Table 2F), where PCs in Chrna2-Cre+ mice exhibited an overall increased activity in the prehension window compared to PCs in control mice (Figure 1J).

We observed that a portion of prehensions overlapped with an already initiated prehension, within the time window of 2 s. Specifically, in the first session, 3.17% of successful and 36.07% of error prehensions occurred within 2s of a succeeding prehension. Similar proportions were found in later sessions: during sessions 2 (6.08% success, 28.26% error), 5 (1.79% success, 28.12% error), and 6 (4.76% success, 29.71% error). When considering all sessions combined, we found a total prehension overlap of 4.0% for successful and 29.91% for error prehensions. Perhaps this result is related to the fact that, when mice fail to get a pellet, they typically try again shortly, resulting in prehension overlaps. To address whether prehension-related neural activity affects baseline signals, we performed FDR-corrected trial-by-trial comparisons of adjacent trials, finding significant effect of the previous trial in the current trial in only 17.7% of pairs, indicating minimal influence of prior trials on calcium activity. Next, we selected trials not preceded by another trial for ≤ 6 s (3x the prehension window size) and compared the activity amplitude between the (−5, −3) and (−3, −1) 2 s time windows relative to prehension onset (Supplemental Figure 3A-H). This analysis revealed no genotype- or session-related differences in baseline amplitude between epochs, though an interaction between accuracy and epoch emerged in the second session (Supplemental Table 2G), driven by lower fluorescence in successful versus failed prehensions for control mice within the (−5, −3) window (Supplemental Table 2H; Supplemental Figure 3I). Collectively, these results confirm that prehension-related neural activity does not systematically alter non-prehension epochs.

We next aligned cell activity to prehension onset (t = 0), and measured the average PC population activity across two distinct phases: a one-second window before prehension (motor planning) and another one-second window during prehension (motor execution). We found similar effects (Supplemental Table 2I-J; Supplemental Figure 2J); reduced PC amplitudes for successful compared to failed prehensions from the second session in control mice. Also, increased excitation of M*α*2 cells in the sixth session resulted in larger PC amplitudes when compared to controls (see summary Figure 6A). While baseline PC activity was decreased in Chrna2-Cre+ mice, the amplitude of prehension-related PC activity was actually increased in these animals, particularly in session 2, which suggests a complex relationship between M*α*2 cell excitability and motor performance.

Finally, we investigated whether the activity proportion between planning and execution phases is affected by increased M*α*2 cell excitability, training, or prehension accuracy. We quantified the ratio of activity between the execution (during) and planning (before) epochs to assess how these factors modulate this proportion. We found a three-way interaction effect of genotype, session, and prehension accuracy (Supplemental Table 2K; Supplemental Figure 4), with genotype effects for both accuracies on the first session and successful prehensions during the second session (Supplemental Table 2L-N), and accuracy effects in control mice across the second and fifth sessions (Supplemental Table 2O-P). In control mice, the balance of PC activity progressively shifted toward movement execution for successful prehensions as the motor skill was refined (sessions 1–5). Following the pellet-position change (session 6), this trend reversed, with activity shifting back toward the planning phase during successful prehensions. This suggests that cortical dynamics are reconfigured during movement refinement. However, Chrna2-Cre+ mice exhibited a markedly reduced reconfiguration of the planning/execution balance across sessions, suggesting that increased Chrna2 Martinotti-cell excitability alters the normal adaptation of cortical activity patterns.

### Increased M*α*2 cell excitation resulted in earlier PC firing during retraining

Changes in sequential activation of PCs are linked to learning [64]. Therefore, we next analyzed the temporal profiles of PCs, and identified the time point where they reached their maximum activity within the prehension window (Figure 2A-B). Using the same dataset as above (n = 6 mice; 3 Control, total of 1070 cells for 4 sessions, average of 105.2, 102.8 and 59.5 for each animal; 3 Chrna2-Cre+, total of 403 cells for 4 sessions, average of 45, 53.8 and 16), we compared the response peak latency between genotypes, sessions, and prehension accuracy while separating neurons based on the prehension phase where the activity peak is reached. For each session, the activity of each PC was averaged and normalized within the prehension window around prehension onset. We sorted individual neurons based on their latency to peak during successful prehensions and enforced the same sorting order for failed prehensions (Figure 2C). The peak latency analysis showed that PC temporal patterns were strongly affected by prehension accuracy (Supplemental Table 3A-C; Supplemental Figure 5; Figure 2D). Many neurons exhibited shifts in their peak activity timing during failed prehensions relative to successful prehensions, such that neurons classified as “before neurons” in the successful prehensions could peak during execution in failed trials, while some execution-related PCs (“during neurons”) could shift their peak activity to before movement onset. These findings indicate a reorganization of the temporal recruitment pattern of PCs during failed prehensions. In control mice, training progressively increased the temporal separation between “before” and “during” execution neurons during successful prehensions, with “before-neurons” peaking earlier and “during-neurons” peaking later (Figure 2D, left). This temporal organization was disrupted during failed prehensions and was altered by increased M*α*2 cell excitability, which produced delayed responses during session 2 and premature responses during task retraining in session 6 (Supplemental Table 3D-E; Figure 6A).

**Figure 2.**
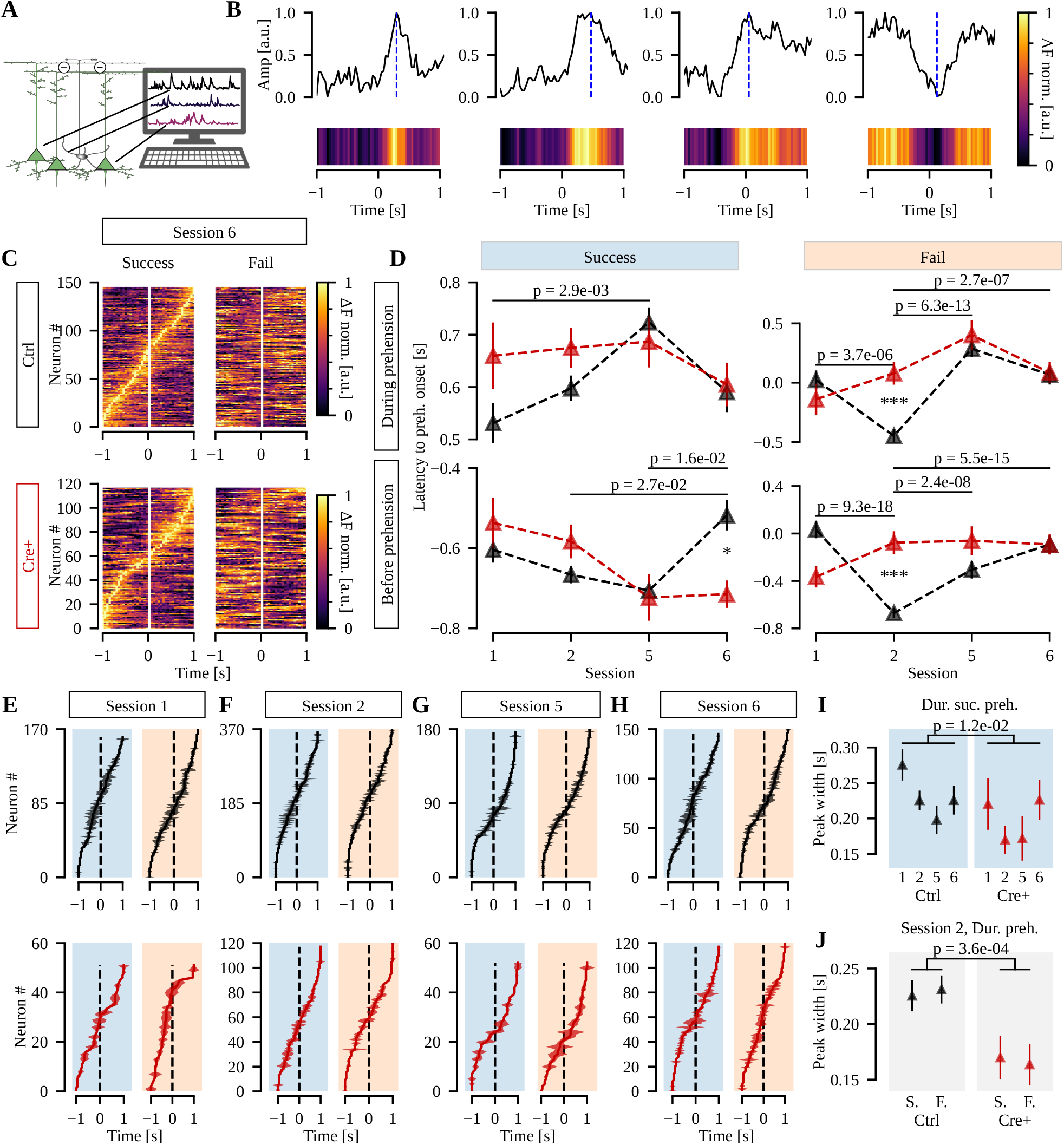
Increased excitation of M*α*2 cells is associated with delayed PC activity during motor performance refinement and premature PC activity on task retraining. A) Illustration of recorded calcium signals from GCaMP6f-expressing PCs. B) Representative examples of average fluorescence traces around prehension onset for four different neurons, showing their normalized fluorescence curve (ΔF detr. over time) represented in a colormap and latency of detected activity peaks (dashed blue line). C) Normalized average fluorescence matrices (Neuron x Time x F) for successful and failed prehensions, for control (top) and Chrna2-Cre+ (bottom) mice, showing temporal activation patterns for each neuron on session 6. Neurons were sorted according to their temporal profile for successful prehensions, and follow the same order for failed prehensions. D) PC peak latency for both genotypes at each session before (bottom) or during (top) successful (left) and failed (right) prehensions. All significance bars refer to significant pairwise differences between sessions for controls; and asterisks represent significant pairwise differences between genotypes for a particular session (Scheirer-Ray-Hare followed by Mann-Whitney U). E-H) Peak latency with a shaded area indicating the peak width during sessions 1, 2, 5 and 6 for each neuron. At each plot, neurons were sorted according to their temporal profile. I-J) PCs response peak width during successful prehensions separated by genotype and session (I), and during prehensions from session 2 separated by genotype and prehension accuracy (J). S., successful; F., failed; B., before; D., during prehension. n cells per session = 162, 363, 399, 146 for Controls; 106, 126, 53, 118 for Chrna2-Cre+ (Scheirer-Ray-Hare).

We also measured the width of the response peak, a measure of the temporal specificity of cell activation that is proposed to influence motor accuracy [65, 66]. Broader response peaks indicate less temporal specificity. We analyzed the response peak width for each genotype across all training sessions, separating prehensions by accuracy and phase according to peak latency (as done for the latency analysis). For visualization purposes only, neurons were resorted for each accuracy, to better represent the peak width as a shaded area around the neuron peak latency (Figure 2E-H). The width of PC activity peaks was most strongly influenced by the session followed by genotype (Supplemental Table 4; Supplemental Figure 6; Figure 2I-J), where PCs in Chrna2-Cre+ mice displayed overall sharper peaks compared to PCs in control mice, particularly during session 2 (Figure 6A). One possible mechanism for this is enhanced M*α*2 cell inhibition of PC apical dendrites, which could reduce dendritic calcium spikes and prolonged dendritic depolarization, thereby limiting burst firing and shortening the duration of PC responses [67].

### Increased M*α*2 cell excitation resulted in PC assembly salience rigidity, distribution compaction, and increased resilience

We next hypothesized that examination of PCs Hebbian assemblies could provide insights into how increased excitation of M*α*2 cells modulate cortical processing. In the assembly analysis, we focused on sessions 1, 2 and 6, in which PC identity could be matched across sessions. On average, 84.8 ± 14.2 PCs were kept for the analysis (n = 4 mice; 2 Control, total of 70 assemblies for 3 sessions, average of 14 and 9.3 for each animal; 2 Chrna2-Cre+, total of 49 assemblies for 3 sessions, average of 11.7 and 4.7). Prehension accuracy did not influence assembly identification, since we based it on neuronal activity across the 10-minute session recording, rather than only during the prehension window (Figure 3A). Assemblies consisting of less than four neurons, and neurons that were not part of any assembly, were omitted from the analysis. In the first session, we identified a total of 23 assemblies in controls and 19 assemblies in Chrna2-Cre+ mice, with an average number of 5.6 ± 0.4 cells in each assembly for each session. The number of neurons within each assembly remained fairly consistent during sessions 2 and 6 (Figure 3B-E).

**Figure 3.**
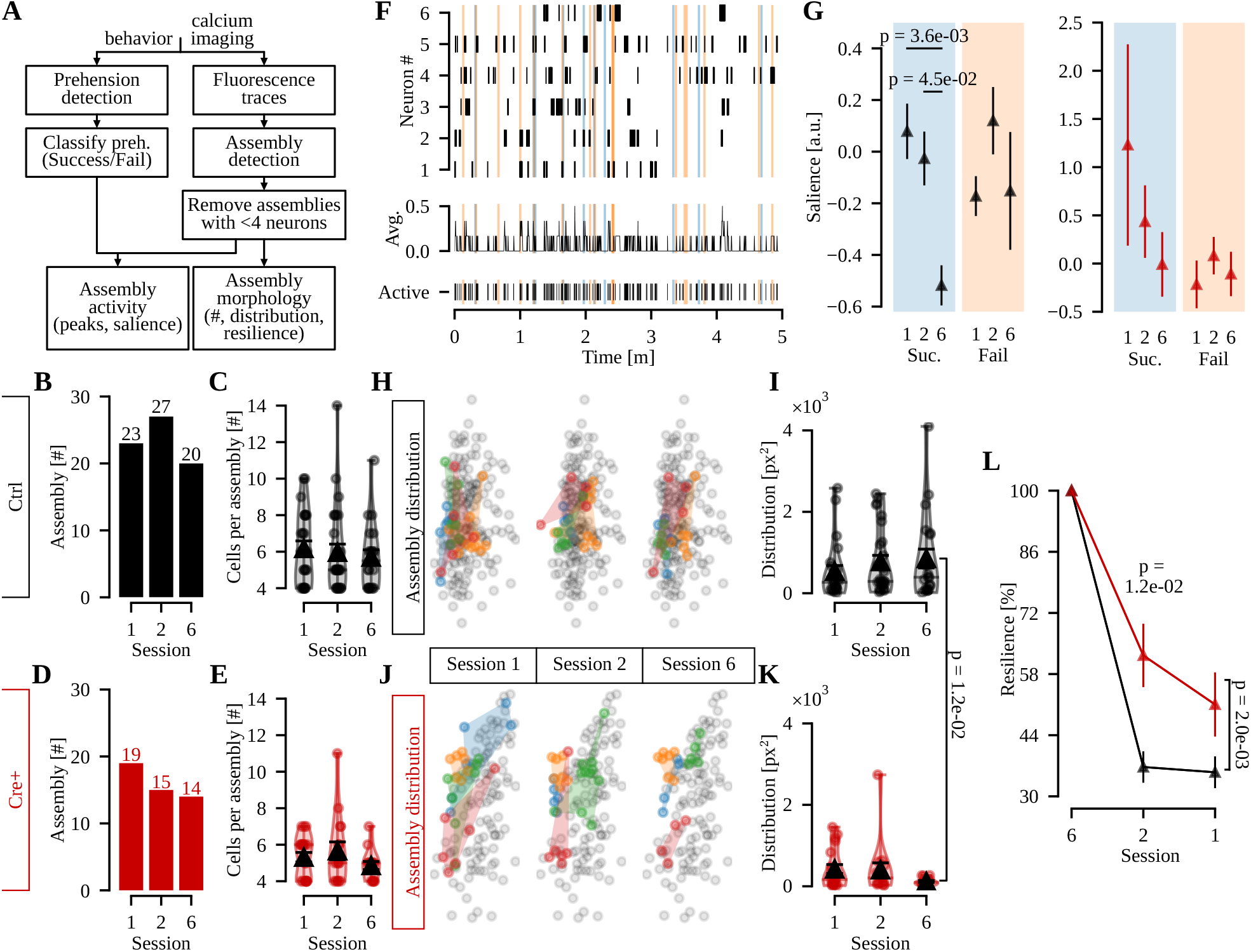
Mice with increased M*α*2 cell excitability display assemblies with less prehension-related activity, decreased spatial area and increased resilience. A) Flowchart of the analysis pipeline. B-C) Number of assemblies and number of cells per assembly for control mice, detected at each session. D-E) Same as B-C, but for Cre+ animals. F) Activity of one representative assembly. Top, rasterized activity over the full recording time of the six neurons forming the assembly. Mid, the average of neuron activity. Bottom, the estimated rasterized activity of the assembly as a unit. Blue and orange shades show the successful and failed prehension windows, respectively. G) Salience of assemblies during the whole 2s window for each session, grouped by genotype and prehension accuracy (Mann-Whitney U). H-I) Representative examples of 4 color-coded assemblies detected at session 6 and traced back to session 1 (H), and spatial distribution of detected assemblies at each session for control mice (I). J-K) Same as H-I, but for Cre+ animals (Mann-Whitney U). L) Resilience of assemblies for each genotype, for assemblies detected at session 6 and followed back to session 1 (ANOVA followed by t-test). Suc., successful; Fail, failed.

Motor training is associated with a selective increase in connectivity among task-related PCs [68]. On a PC population level, we examined assembly salience, here defined as the activity of the assembly within the prehension window in relation to its own activity outside the prehension window (ranging from −100 to 100; Figure 3F). The salience data for single assemblies varied from −2.04 to 24.62 and we found an overall effect of the training session on assembly salience (Supplemental Table 5A). Assembly activity during successful prehensions in control mice was less salient in session 6 compared to sessions 1 and 2 (Supplemental Table 5B-C; Figure 3G and 6B). We found no other differences in salience for failed prehensions or between training sessions in Chrna2-Cre+ mice. This analysis suggests that the decreased activity observed in single neurons from control mice also occurs at the assembly level, where activity within the prehension window is decreased in relation to assembly activity outside the prehension window.

Functional imaging studies in humans show that brain activity varies regionally during different stages of motor learning [69]. Hence, we plotted the position of the individual PCs in each assembly and measured the spatial distribution which each assembly covered in the cortex. We found smaller assemblies in Chrna2-Cre+ mice compared to control mice in session 6 (Supplemental Table 5D-E; Figure 3H-K), suggesting that upon increased M*α*2 cell excitation assemblies were composed of PCs spatially closer together (Figure 6A-B). To measure PC assembly plasticity, we investigated how much the assemblies had to change their neuronal composition to reach their final configuration in session 6, quantified here as assembly resilience. Resilience was calculated taking the assembly composition reported as the percentage of neuron match between sessions, with 100% being an assembly that did not incorporate or remove any neurons over time. We found that genotype and training session affected assembly resilience (Supplemental Table 5F), where assemblies in Chrna2-Cre+ mice displayed increased resilience compared to assemblies in control mice in session 2 (n = 20 assemblies control and 14 assemblies Chrna2-Cre+ traced back from session 6; Supplemental Table 5G; Figure 3L). Together, these results suggest that PC assemblies undergo less modifications on their neuronal composition upon increased M*α*2 cell excitation (Figure 6A-B).

### Increased M*α*2 cell excitation resulted in improved execution of a learned motor task

So far, we found that increased M*α*2 cell excitation resulted in several effects on PC population and assembly plasticity in the motor cortex (Figure 6A-B). Notably, despite these changes in neural activity, we found no effect of genotype on prehension performance (Figure 1F-H; Supplemental Table 1A-C), which highlights a potential dissociation between changes in neural activity and performance outcomes. Since we observed changes related to plasticity with no effect on performance outcome when activating M*α*2 cells repetitively, we instead wanted to evaluate the potential effect of M*α*2 cell activation on already acquired motor skills. To investigate the possible influence of increased M*α*2 cell excitability on already learned motor skills, we trained mice (Chrna2-Cre+, n = 5; control, n = 6) for seven sessions prior to injecting them with hM3Dq and then evaluated their performance in a prehension task (Supplemental Figure 7A). In the first three post-injection sessions (8, 9 and 10), saline was administered to assess potential injection effects, while CLZ was administered in the following three sessions (11, 12 and 13) to measure the potential impact (Figure 4A). We found an overall effect of session in the success rate (Supplemental Table 1G-H), and when grouping by genotype we found that CLZ treatment increased the success rate in Chrna2-Cre+ mice compared to saline treatment, which was not observed in control mice (Supplemental Table 1I-L; Figure 4B). These results suggest that enhancing M*α*2 cell excitability improved the performance of a motor task relying on an already refined skill (Figure 6C).

**Figure 4.**
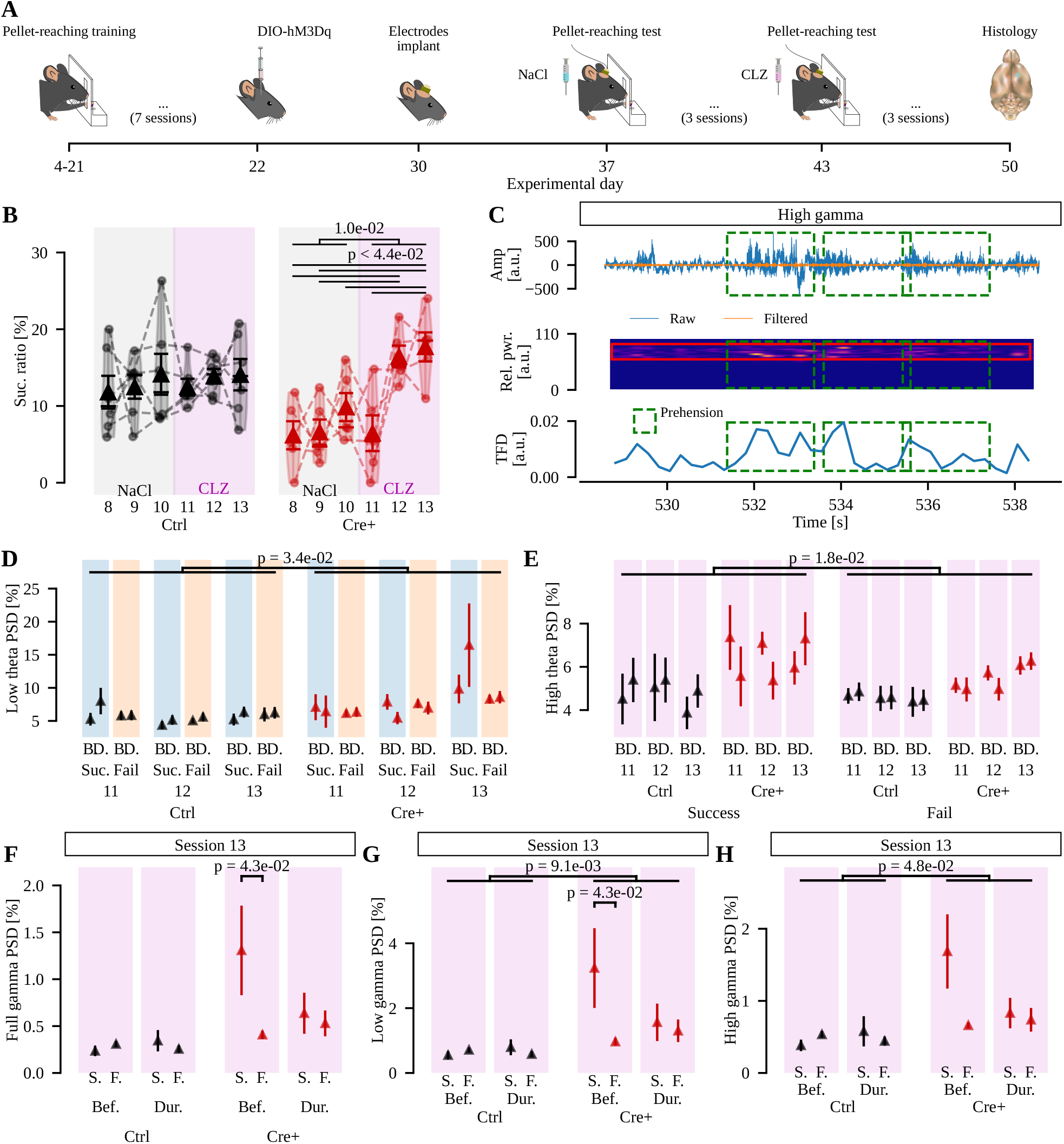
Mice with increased M*α*2 cell excitability display increased prehension success rate, increased LFP low theta and gamma power. A) Timeline of the experiments. Animals were trained to reach for the pellet and after the 7th session, the hM3Dq was injected and the LFP electrodes implanted. B) Prehension success ratio for control (Left, black) and Chrna2-Cre+ (Right, red) mice in the sessions with saline and clozapine injection (gray and purple shades, respectively; ANOVA followed by t-test). C) Example of time-frequency domain power processing, with three prehensions highlighted (green dashed squares). Top, raw (blue) and high gamma-filtered (orange) LFP traces. Mid, spectrogram from the filtered signal, with highlights for the high gamma band (red square). Bottom, the time-frequency domain power over time, calculated as the average of the previously highlighted spectrogram region. D-H) Relative power for low theta (ANOVA), high theta (ANOVA), full gamma (ANOVA-type), low gamma (ANOVA-type) and high gamma (ANOVA), respectively. D and E shows the full dataset due to the observed overall effects, while for F-H only isolated effects for session 13 were found. Shades: blue and orange (successful and failed prehensions); magenta (sessions under CZP treatment). Abbreviations: B. or Bef., before prehensions; D. or Dur., during prehensions; S. or Suc., successful prehensions; F. or Fail, failed prehensions. n mice = 6 Controls and 5 Chrna2-Cre+.

Low and high sub-bands within theta and gamma frequencies are associated with different functional roles in attentional control, sensory integration, and fine motor control. To gain deeper insight into how M*α*2 cell modulation affects motor planning and execution, we recorded local field potentials (LFPs) in the same mice during task performance in the CLZ-treated sessions (sessions 11–13). This allowed us to correlate neural activity patterns with specific phases of the task. We separated successful and failed prehensions and analyzed one-second time windows before (planning phase) and during (execution phase) task onset. Using spectrograms, we calculated power in various frequency bands across the LFP signal, spanning 1–100 Hz, with specific focus on the theta (4–12 Hz) and gamma (30–90 Hz) ranges. The theta band was further divided into low (4–7 Hz) and high (8–12 Hz) frequencies, and gamma was divided into low (30–40 Hz) and high (60–90 Hz) frequencies (Figure 4C).

LFP analysis provided a detailed view of M*α*2 cell involvement, showing increased power in the full theta band and the low theta band in Chrna2-Cre+ mice compared to controls (Supplemental Table 6A-C; Figure 4D). Notably, in the high theta band, we observed higher power in successful than failed prehensions (Supplemental Table 6D; Figure 4E). For the gamma band, we only found significant effects on session 13, which showed similar trends as high theta, with higher power in successful than failed prehensions across gamma bands (Supplemental Table 6E-F; Figure 4F–H). For the low gamma band, Chrna2-Cre+ mice displayed higher power than controls, especially in successful prehensions (Supplemental Table 6G-H; Figure 4G). High gamma power was also elevated in Chrna2-Cre+ mice compared to controls (Supplemental Table 6I-J; Figure 4H).

We next evaluated the LFP power outside prehension epochs, i.e., when no prehension movement was being performed. Analysis showed an interaction effect of genotype and session across all frequency bands (Supplemental Table 6K). For full, theta, and beta bands, these interactions were driven by a session effect in Chrna2-Cre+ mice (Supplemental Table 6L, Supplemental Figure 7B-E), with LFP power in session 13 consistently decreased compared to previous sessions, not observed in control mice. When subdividing theta and gamma into low and high subdivisions, low-theta power was affected both by genotype and training session (Supplemental Table 6M-N, Supplemental Figure 7F). Similarly, genotype affected both low- and high-gamma bands (Supplemental Table 6O-P, Supplemental Figure 7G-H). No effects were found for the high-theta band.

Collectively, these findings suggest that M*α*2 cell activation plays a role in increasing low theta and gamma power during motor planning and execution phases of an already refined skill, while reducing overall LFP power when no prehension is being performed, supporting a role for M*α*2 cells in fine-tuning motor performance (Figure 6C).

### Ablation of M*α*2 cells affected fine motor functions

To investigate how the absence of M*α*2 cells might affect motor execution, we used a Caspase-based virus approach to selectively ablate these cells in the forelimb area of the M1 of Chrna2-Cre/tdTomato mice. After unilateral injections (n = 2; Figure 5A), the deletion resulted in a reduction of 80.4% of the number of tdTomato-positive cells on the injected side compared to the uninjected side within the same mouse (Figure 5B). Initially, we used a pasta handling test to assess forepaw dexterity that does not require training and found that bilaterally injected Chrna2-Cre+ mice dropped more pasta than control mice (n = 3 Chrna2-Cre+, 6 controls; Supplemental Table 1M), although their handling times were similar (Figure 5C-E). We also used a hanging wire test to evaluate grip strength and endurance [70], but found no differences between the two groups in terms of number of falls or time spent on the wire (Figure 5F-H). In the prehension task, we found an overall effect of training session in the success ratio (Supplemental Table 1N), but we did not find an effect of genotype in the prehension’s success ratio (Supplemental Table 1O), only a punctual pairwise difference between sessions 7 and 10 (Figure 5I, Supplemental Table 1P). Thus, ablation of M*α*2 cells did not affect prehension task success, but reduced dexterity in the pasta handling test. This suggests that M*α*2 cells may have a role in orchestrating precise forelimb movements that do not require previous specific training (Figure 6C). Together with results from the group where mice with increased excitability of M*α*2 cells displayed improved performance compared to baseline, these findings suggest that although M*α*2 cells may contribute to fine-tuning motor performance, they are not essential for executing refined complex movements.

**Figure 5.**
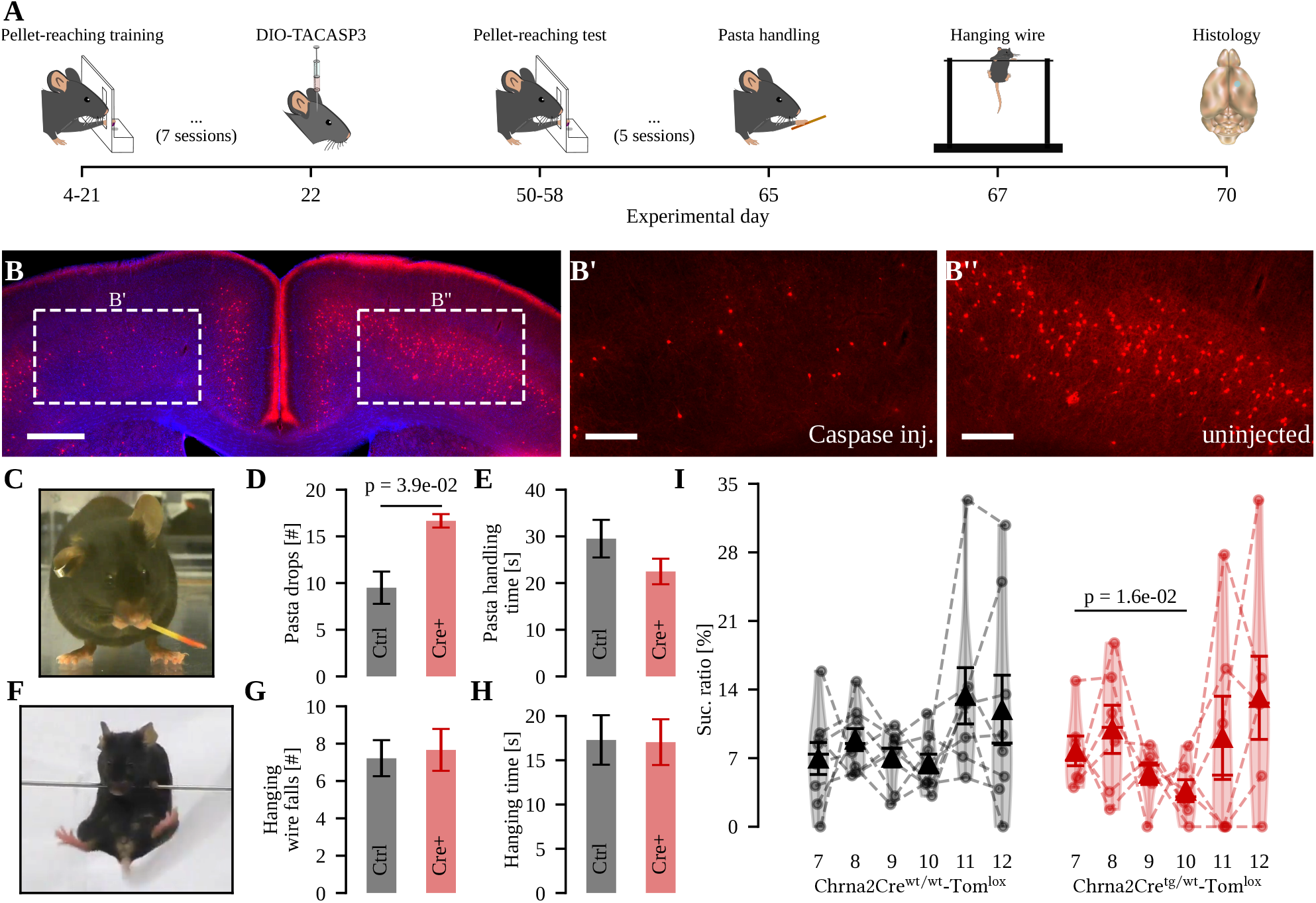
Apoptosis in M*α*2 cells decreased pasta handling performance. A) Timeline of the experiments. B) Unilateral injection of Cre-dependent TACASP3 decreased the amount of Chrna2-Cre+ cells in the motor cortex by 80.4%. Enlargement of the injected motor cortex (B’), and contralateral non-injected side (B”). Scalebars: 500µm (B) and 200µm (B’ and B”). C) Animal performing the pasta handling task. D-E) Graph of results on pasta drops and time spent in the task (ANOVA). F) Animal performing the hanging wire task. G-H) Graph of results on falls and time spent in the task (ANOVA). n mice = 6 Controls and 3 Chrna2-Cre+. I) Single pellet prehension task before (session 7) and after ablation (sessions 8-12) for WT (black) and Chrna2-Cre+ mice (red). n mice = 8 Controls and 6 Chrna2-Cre+ (ANOVA).

## Discussion

Here we investigated how M*α*2 cells impact PC population dynamics during motor performance refinement and execution. We found that, upon increased M*α*2 cell excitation, parameters of plasticity at the cell, population and assembly level, all indicate decreased plasticity (Figure 6A-B). Whereas increased M*α*2 cell excitation during training did not affect prehension accuracy, it did improve the success rate when chemogenetic activation was initiated in mice that had already refined the movement performance (Figure 6C). Thus, our data suggest that increased M*α*2 cell excitation seems to reduce PC plasticity while possibly facilitating an already acquired motor skills.

**Figure 6.**
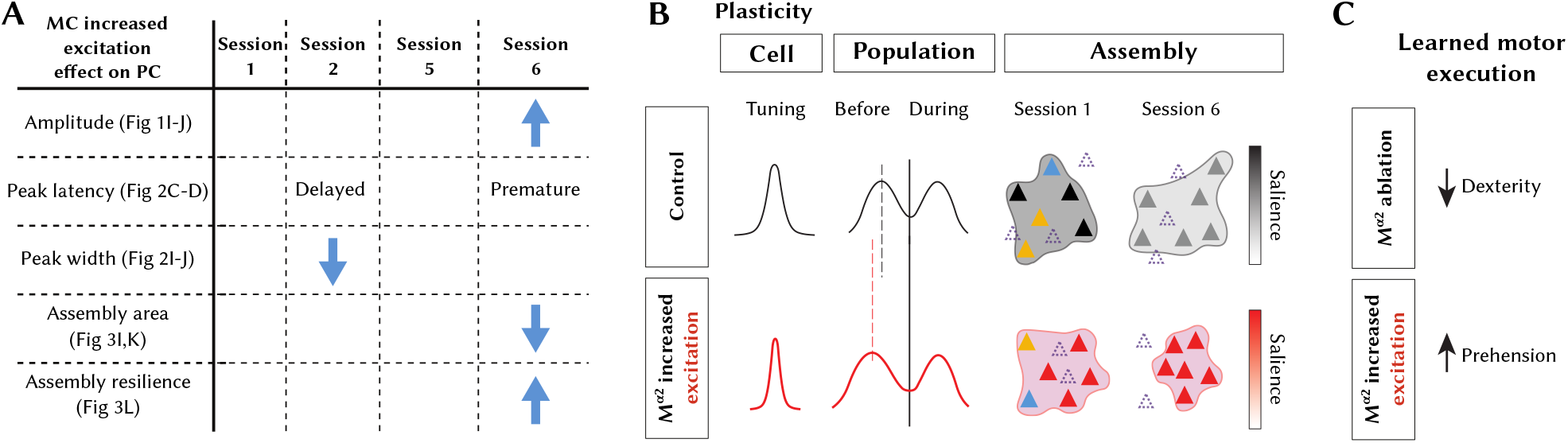
Summary of findings. A) Table summarizing the effects of increased M*α*2 cells excitation on PCs. B) Schematic outline illustrating plasticity parameters. Increased excitation of M*α*2 cells resulted in more narrow tuning of layer 5 PCs, a temporal shift of the average population activity both in the motor planning and execution phases, increased resilience, reduced physical assembly coverage and rigid salience upon retraining. Dashed triangles mark PCs that are part of the retrained assembly but not the naive assembly, and vice versa. Blue and yellow depict PCs in the naive assembly that are part of another assembly in the retraining session. Salience values are represented by the degree of color in PCs and assembly distributions. C) M*α*2 cells influence fine motor movements, where caspase induced ablation of M*α*2 cells resulted in decreased dexterity, whereas increased excitation of M*α*2 cells resulted in increased accuracy.

Motor cortex activity during skilled behaviors such as prehension does not reflect event-by-event correlations between behavioral actions and single-neuron calcium transients. Instead, motor output emerges from distributed population dynamics in which individual neurons contribute sparsely and variably across trials. As demonstrated in classical studies of corticospinal function [71], there is no direct or necessary link between single-unit firing and specific movements; neural activity must be analyzed at population level to reveal consistent relationships with behavior. These population-level dynamics are further constrained by the inherent limitations of calcium imaging, where signals are temporally filtered and act as nonlinear proxies for spiking activity, showing low sensitivity to isolated spikes and a strong bias toward bursts or clustered firing patterns. Consequently, action potentials, particularly those occurring in isolation, may fail to generate detectable transients. The observed discrepancy between prehension events and calcium events thus reflects fundamental principles of motor cortex coding and technical constraints of imaging, rather than a failure to capture relevant neural dynamics. Our analysis therefore focuses on population-level metrics such as assembly activity patterns, trial-averaged responses, and dimensionality reduction, which provide a more accurate representation of how cortical circuits support skilled movement.

Lateral inhibition, a canonical function of inhibitory circuits and Martinotti cells (MCs) in particular, sharpens excitatory responses and promotes competitive interactions among pyramidal cells [PCs; 72, 15]. By suppressing neighboring PCs, early activated cells can dominate network output, implementing a winner-take-all strategy in which temporally leading PCs emerge as influential “leader” cells. Consistent with this framework, motor learning shifts the temporal activation of L2/3 PCs toward earlier firing [11], and our calcium imaging data likewise suggest that temporal activation patterns, rather than response amplitudes, predict successful prehension. Successful trials were characterized by an early PC peak during the planning phase followed by a later peak during movement execution, underscoring the importance of precisely structured temporal dynamics.

Although SST interneurons have been shown to enable and maintain sequential PC activation during motor learning [11], accumulating transcriptomic and circuit-level evidence demonstrates that SST cells comprise multiple molecularly and functionally distinct subtypes [73, 21], which can display layer-specific and even opposing activity patterns [16]. Broad SST manipulations therefore likely engage heterogeneous populations with divergent functions. In contrast, we targeted a homogeneous layer 5 Martinotti subtype [M*α*2 cells; 24] and found that increasing M*α*2 cell excitability shifted PC peak activity in relation to the movement onset, enhanced directional tuning precision, and improved execution of an already learned movement without affecting acquisition. This contrasts with reports of impaired motor learning following global SST activation [74, 75] and supports the idea that distinct SST subpopulations differentially regulate learning and execution phases. Mechanistically, enhanced M*α*2 cell activity likely sharpens tuning [65, 66], promotes temporal synchronization of layer 5 PCs [24], and stabilizes assembly structure during retraining, thereby biasing the emergence of temporally leading “leader” PCs. Together, our findings support a model in which layer-specific SST subtypes sculpt PC temporal dynamics and assembly plasticity in a phase-dependent manner during motor behavior.

Importantly, our conclusion that M*α*2 cell activation dissociates neural dynamics from performance outcomes should be interpreted within the limits of our behavioral measurements. Motor performance was quantified primarily as prehension success rate. While this metric captures task outcome, it does not resolve finer kinematic parameters such as reach trajectory, movement speed, acceleration profiles, or grip force. It therefore remains possible that M*α*2 cell modulation affects additional aspects of motor execution that were not measured here. Future studies combining neural recordings with detailed kinematic analyses will be important to determine whether the observed changes in PC temporal dynamics are also reflected in subtler alterations of movement structure. Temporal properties differ between interneuron populations, and similarly, PCs are likely to be variably inhibited depending on timing, synaptic weights, and the position of the interneuron contact [76]. Notably, a hypothesis postulates a central role of local inhibitory drive in tuning motor circuits for accuracy [65, 66]. A strong inhibitory drive would narrow directional tuning widths and lead to more accurate movements. Our experimental data on motor execution adhere to and support this theory, since we found that heightened excitation of M*α*2 cells, i.e. more inhibition, resulted in improved accuracy and narrower PC activity tuning (Figure 6).

In addition to sharpening responses via lateral inhibition, M*α*2 cells can actively promote synchronization of pyramidal cell (PC) activity. Because each M*α*2 interneuron targets the distal dendrites of multiple PCs, their coordinated firing, or release from inhibition, can impose a shared inhibitory timing signal across a local PC population. This can align the timing of PC spikes and thereby promote synchronous firing. Indeed, in layer 5 neocortex, optogenetic activation of M*α*2 cells were shown to transform previously uncorrelated PCs into temporally synchronized ensembles, particularly among type A pyramidal cells [24]. Such synchronization depends critically on the firing pattern of the interneurons, underscoring that inhibition does not merely suppress activity but actively structures the temporal coordination of excitatory networks. In the context of our data, this provides a mechanistic framework for how increased M*α*2 cell excitation could bias the emergence of “leader” PCs and constrain assembly reconfiguration during retraining by enforcing tighter temporal coupling among layer 5 PCs.

Hebbian learning postulates that when neurons fire synchronously, they reinforce synaptic efficiency and form an assembly. The original theory has been extended to include all cells with above-chance interactions, leading to variable definitions of neural assemblies [77]. We here adhere to the proposition that assemblies are formed by neurons that together increase their average firing rates for a specific period [78, 39], and that they can harbor varying degrees of temporal precision, scale and internal structure [79]. According to the extended theory, assemblies dynamically form, reconfigure, and fire simultaneously, with a reduced number of cells triggering the entire assembly [80]. The theory also proposes that one single neuron could participate in multiple neural assemblies, leading to distributed information coding. When PCs in our data set were sorted into assemblies, we found that the average number of cells composing the assemblies were similar regardless of M*α*2 cell activation state. Notably though, the distribution of each assembly, which remained constant during sessions 2 and 6 in control mice, were smaller during session 6 in mice with increased M*α*2 cell activity, demonstrating that M*α*2 cell activity can affect PC cell assembly internal structure. In sessions 2 and 6, increased M*α*2 cell excitation resulted in an increased resilience in PC assemblies, suggesting that increased M*α*2 cell excitation decreases PC assembly plasticity. Accordingly, during session 6, the assembly salience decreased in control mice, but not in Chrna2-Cre+ mice, further supporting that increased M*α*2 cell excitation decreases plasticity and network reorganization (Figure 6A-B).

In the motor cortex, different motor tasks induce dendritic calcium spikes on different apical branches of individual PCs, which reduces the chance of acquired skills to be disrupted by the formation of new motor programs. Intriguingly, after SST interneuron ablation, the maintenance of synaptic potentiation acquired during previously learned tasks is lost, resulting in decreased performance [3]. We found, after ablation of M*α*2 cells, an increased number of drops in the pasta handling task. Thus, M*α*2 cells, as part of the SST population, also seem important for facilitating execution and refining already acquired fine motor skills. Further, it has been shown that increased SST interneuron activity helps maintain previously acquired motor skills when executing a slightly different motor task, whereas decreased SST interneuron activity is important for improved motor performance and learning of the initial motor task [11, 81]. Also, the initial learning phase involves weak cortical activation of SST interneurons, and even though individual SST interneurons show diverse activity profiles during movement phases in a lever-press task, the average activity of SST interneurons increases as learning progresses [81]. We found that increased excitation of M*α*2 cells during execution of a learned motor task, the single-pellet prehension task, improved the success rate. However, in contrast to what has been shown for SST interneurons in layer 2/3 [11], repetitive increased excitation of M*α*2 cells during training did not affect motor performance. Hence, it seems that increased excitation of M*α*2 cells, resulting in decreased PC assembly reconfiguration and plasticity, mainly influences the execution of already acquired motor skills and that decreased motor learning during SST interneuron activation is likely mediated by other SST interneuron subpopulations. It has been suggested that layer specific network activity takes place during learning of a motor task, where the L2/3 network seem to represent coordination of signals during learning, whereas L5a may participate in networks for well-learned movements [82].

The network of inhibitory interneurons is essential for generating gamma rhythms [83]. Previous studies have highlighted the critical roles of parvalbumin and somatostatin interneurons in sustaining gamma activity in both the cortex and hippocampus [84, 85]. Research involving local field potentials (LFPs) in monkeys and humans has shown an increase in high gamma power (70–170 Hz) alongside a decrease in beta power (8–30 Hz) prior to and during forelimb movements [86, 87, 88]. In rats, high gamma power was also observed during upward movements in a reaching task, measured through epidural field potentials (EFPs) and LFPs [89], where both demonstrated an increase in high gamma power during upward reaches. Additionally, power increases in the high gamma band (70–170 Hz) were noted just before movement onset for upward, left, and right reaches across different tasks [89]. Our results pinpoint a role for M*α*2 cells in gamma and theta oscillations during motor performance, particularly in the context of successful versus failed trials. The elevated high theta power in the success trials compared to failed trials, and increased power in low and high gamma oscillations in Chrna2-Cre+ mice compared to controls, suggests that M*α*2 cells may be important for motor planning and execution. Overall, these findings reveal a complex interplay between M*α*2 activity, oscillatory brain dynamics, and motor function, prompting further investigation into how these mechanisms impact behavioral outcomes.

A limitation of our design is the absence of a Chrna2-Cre+ group expressing a Cre-dependent control vector without hM3Dq. In principle, we therefore cannot fully exclude effects of viral expression per se. However, several observations argue against this interpretation. Cre-dependent expression alone has repeatedly been shown not to affect behavior or network activity in the absence of DREADD activation [90, 91, 92]. We also controlled for nonspecific clozapine effects by administering the same low dose to Chrna2-Cre+ and control mice, and verified effective hM3Dq activation via c-Fos induction in M*α*2 cells. Finally, we observed no effect of Cre-dependent hM3Dq expression alone in our LFP controls (Figure 4). Together, these data make it highly unlikely that the reported effects are driven by viral expression rather than increased M*α*2 cell activity.

In conclusion, we have identified a role for M*α*2 cells in PC population dynamics, motor skill refinement and execution. We find that activation of layer-5-specific Martinotti cells improves the execution of an already refined skill, whereas skill acquisition is unaffected, functionally separating these two aspects of motor activity. Even so, M*α*2 cells are involved in temporal shaping of PC population dynamics and impact several aspects of assembly plasticity during motor training and retraining. Further studies are warranted to address the activity patterns of M*α*2 cells themselves during motor learning and execution, and also whether modulation of M*α*2 cell activity can affect other types of learning and skills emanating from other cortical areas.

## Author Contributions

Methodology (AV, JW, BC, RL), Data curation (AV, JW, BC, KH), Validation (AV, JW, BC), Software (GN, TM), Formal analysis (TM, AV, KH, GN), Visualization (TM, BC, AV, KH, KK), Original draft preparation (KK, TM, AV, BC), Conceptualization (KK, AV, RL), Writing-Reviewing and Editing (KK, TM, BC, AV).

## Acknowledgments

We thank Uppsala University behavioral facility (UUBF) and Charité for support and reagents, Juan Jiang, UmU, for establishing the prehension task.

## Data Availability Statement

The datasets generated and/or analyzed in the current study are available on request.

