## Supplemental material for "Increased layer 5 Martinotti cell excitation reduces pyramidal cell population plasticity and improves learned motor execution"

Thawann Malfatti[1]*, Anna Velica[1]*, Jéssica Winne[1,2,3], Barbara Ciralli[1,2], Katharina Henriksson[1], George Nascimento[2]#, Richardson Leao[2], Klas Kullander[1]

* equal contribution, # deceased 2022

[1]Department of Immunology, genetics and pathology, Uppsala University, Uppsala, Sweden.

[2]Brain Institute, Federal University of Rio Grande do Norte, Natal/RN, Brazil.

Present address:

[3]Department of Biology, University of Maryland, College Park, U.S.

[4]Department of Clinical Neurosciences, University of Cambridge, CB2 0QQ, United Kingdom.

Klas kullander

### M$\alpha$2 cell excitation can be chemogenetically increased

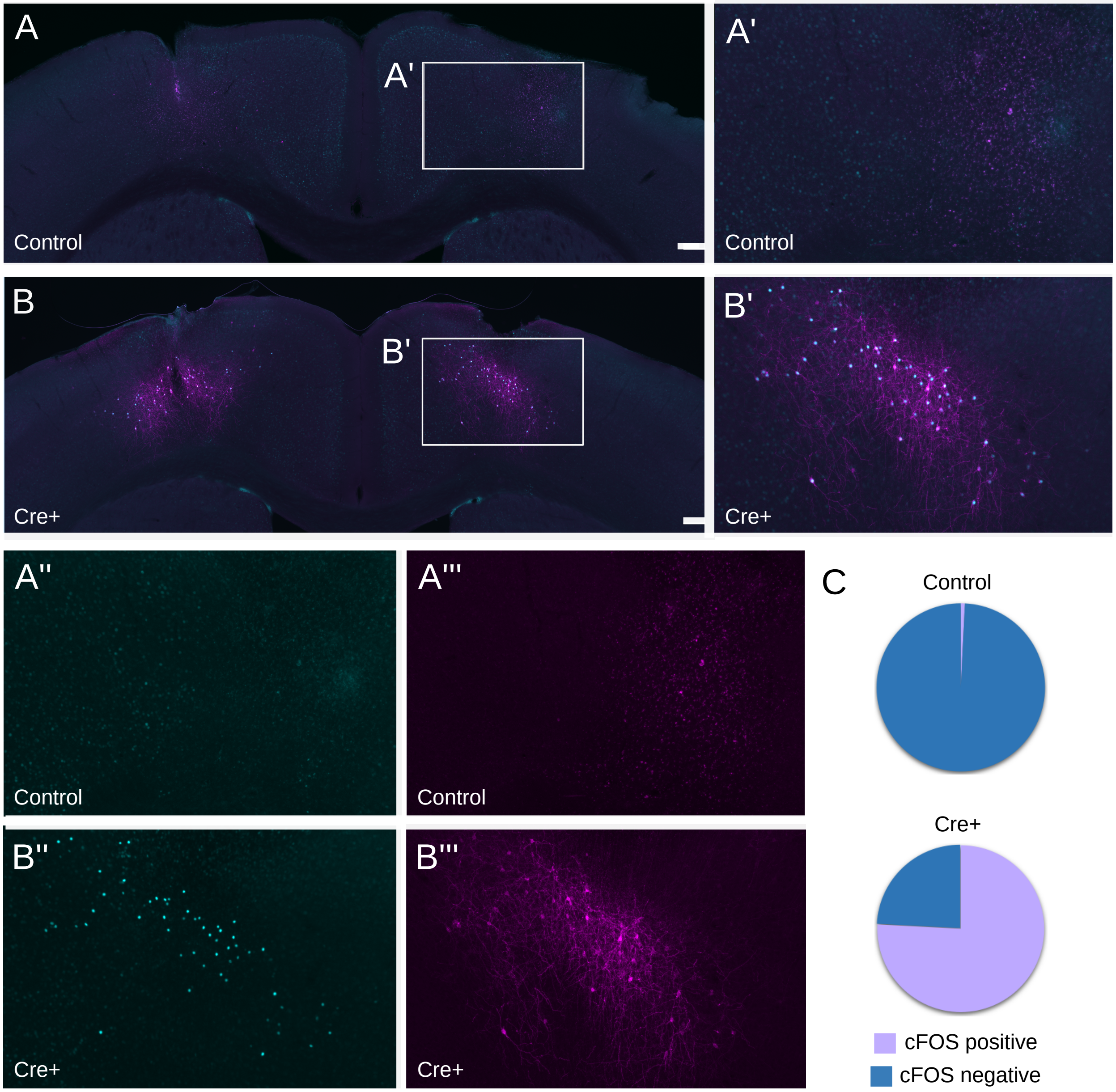


**Supplemental Figure 1:** Validation of the excitatory modified Gq-coupled human M3 muscarinic receptor (hDM3Gq). A-B) Immunolabeling for c-Fos (cyan: A, A', A", B, B' and B") and mCherry (magenta: A, A', A"' and B, B' and B"') after CLZ administration on brain sections from mice injected with the viral vector carrying hDM3Gq and mCherry under Cre-dependent expression. SB 200µm. C) In control mice 0.8% of the mCherry+ cells expressed c-Fos, while in Chrna2-Cre+ mice 76% of the mCherry+ cells expressed c-Fos.

To validate the clozapine (CLZ) induced activation of hM3DGq+ cells, we injected viral vectors carrying the modified Gq-coupled human M3 muscarinic receptor (hDM3Gq) and the fluorescent marker mCherry (Alexander et al., 2009) under Cre-dependent expression and performed immunolabeling for c-Fos, a proto-oncogene expressed in most neurons after depolarization (Morgan et al., 1988, Bullitt 1990; Supplemental Figure 1A-B). In Chrna2-Cre+ mice (n=9, number of sections=346), on average 32 mCherry positive cells per hemisection were found, of which 76% were c-Fos positive. In control mice (n=5, number of sections=60), on average 5 mCherry positive cells per section were found, of which 0.8% were c-Fos positive (Supplemental Figure 1C).

### Prehension-related PC activity decrease with motor training

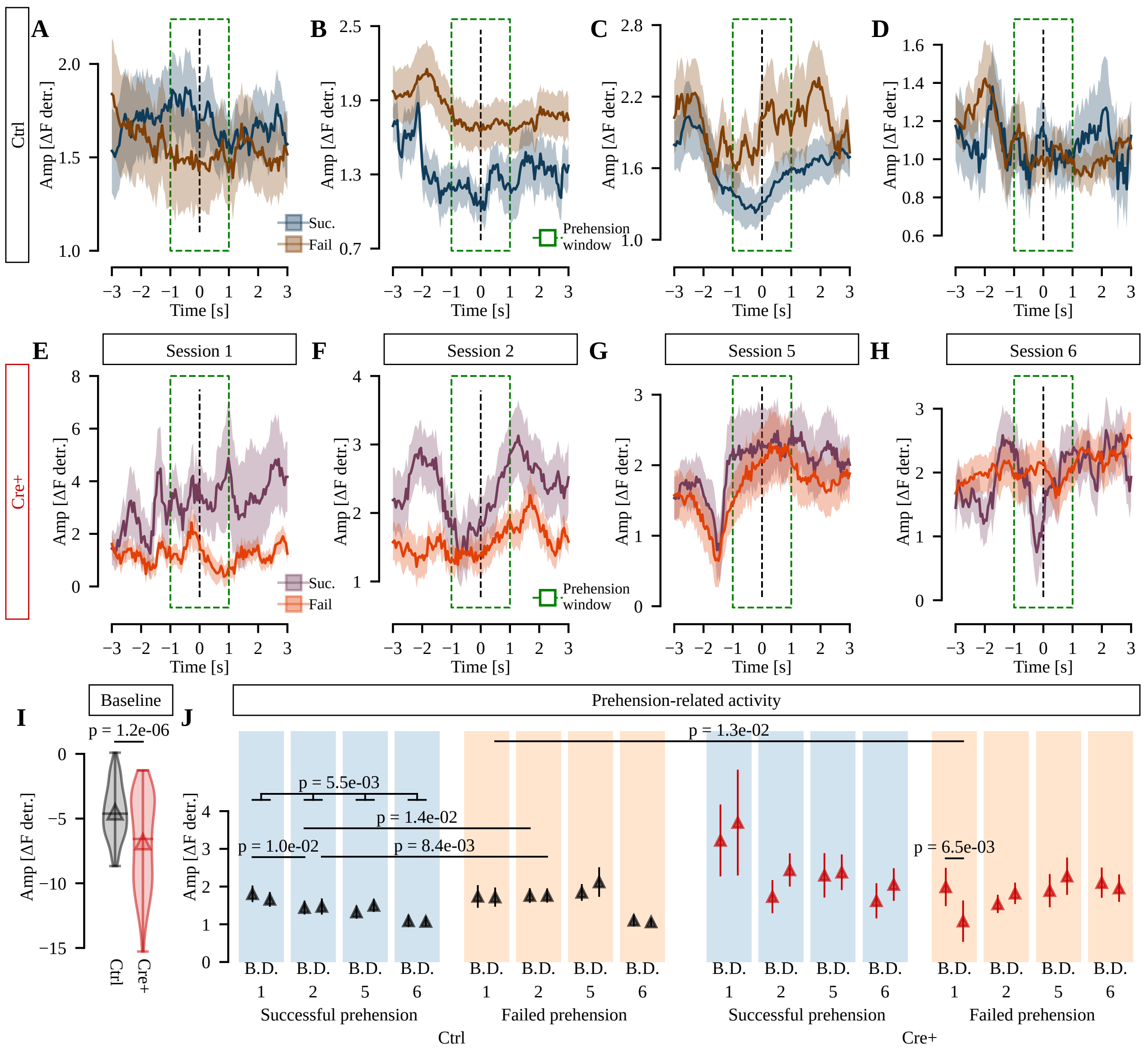


**Supplemental Figure 2:** Prehension-related PC activity decreases for successful prehensions with training in Control mice. A-H) Average traces (lines) and SEM (shades) of prehension-related activity separated by genotype, session and prehension accuracy, including all trials and all neurons. I) Amplitude of PC baseline activity separated by genotype (control, Ctrl, black; Cre+, red). J) Average amplitude of PC activity represented as mean ± SEM, separated by genotype (control, Ctrl, black; Cre+, red), prehension accuracy (successful prehension, blue shade; failed prehension, orange shade), session (naive, N.; learning, L.; training, T.; retraining, R.), and prehension phase (before prehension, B.; during prehension, D.).

The prehension-related activity varies according to genotype, session and prehension accuracy (Supplemental Figure 2A-H). As expected from chemogenetically increasing the excitability of inhibitory interneurons, enhancing M$\alpha$2 cell excitability decreased baseline PC activity in Chrna2-Cre+ mice relative to controls (Supplemental Figure 2I). Specifically, for control mice, prehension-related PC activity both before and during prehension was higher for failed prehensions compared to successful prehensions in the learning session (Mann-Whitney U eff. size $>$ 0.143, p $<$ 1.4e-02; Supplemental Figure 2J). Also, PC activity decreases on successful prehensions as the training progresses (ANOVA-type, F1-LD-F1 model F(3,inf) = 5.37, p = 5.5e-03), especially before successful prehensions in the learning session compared to the naive session (eff. size = 0.204, p = 1.3e-02; Supplemental Figure 2J). These findings suggest that a decrease in PC activity amplitude might be associated with improvement in the task. No such effects were observed for Chrna2-Cre+ mice, however, PC activity during failed prehensions was decreased compared to activity before failed prehensions in the naive session (Wilcoxon signed-rank eff. size = 0.344, p = 6.5e-03; Supplemental Figure 2J), indicating that for naive mice an increased excitation of M$\alpha$2 cells might lead to a changed PC activation pattern that is compensated for with training.

### Prehension-related activity does not affect baseline

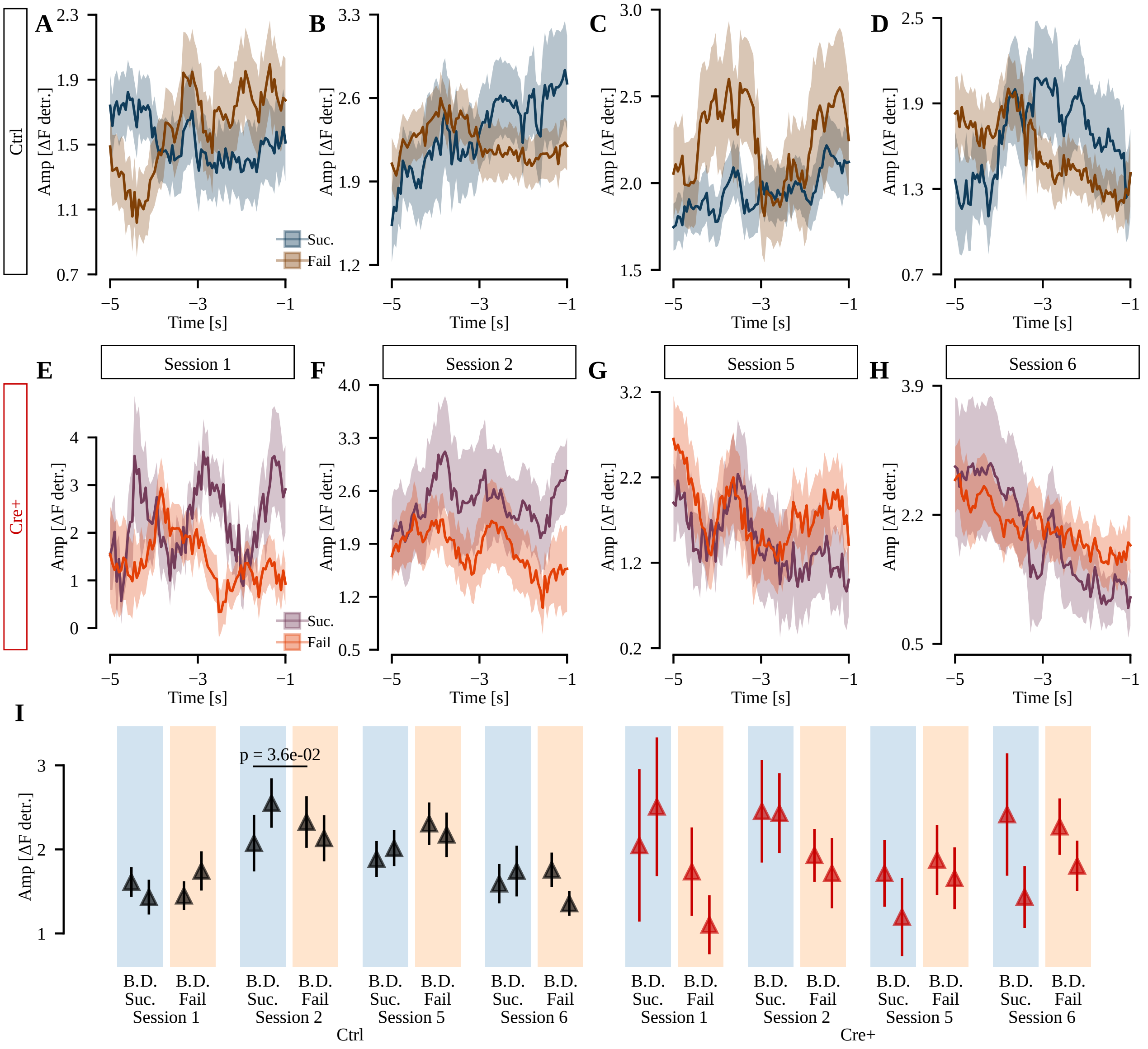


**Supplemental Figure 3:** Baseline is not affected by prehension-related activity. A-H) Baseline average traces (lines) and SEM (shades) -5 to -1s before prehension, separated by genotype, session and prehension accuracy, including all trials and all neurons. I) Average amplitude for baseline fluorescence, separated by genotype, session, prehension accuracy and epoch (-5 to -3s vs -3 to -1s).

### M$\alpha$2 cell excitation can change planning/execution activity balance

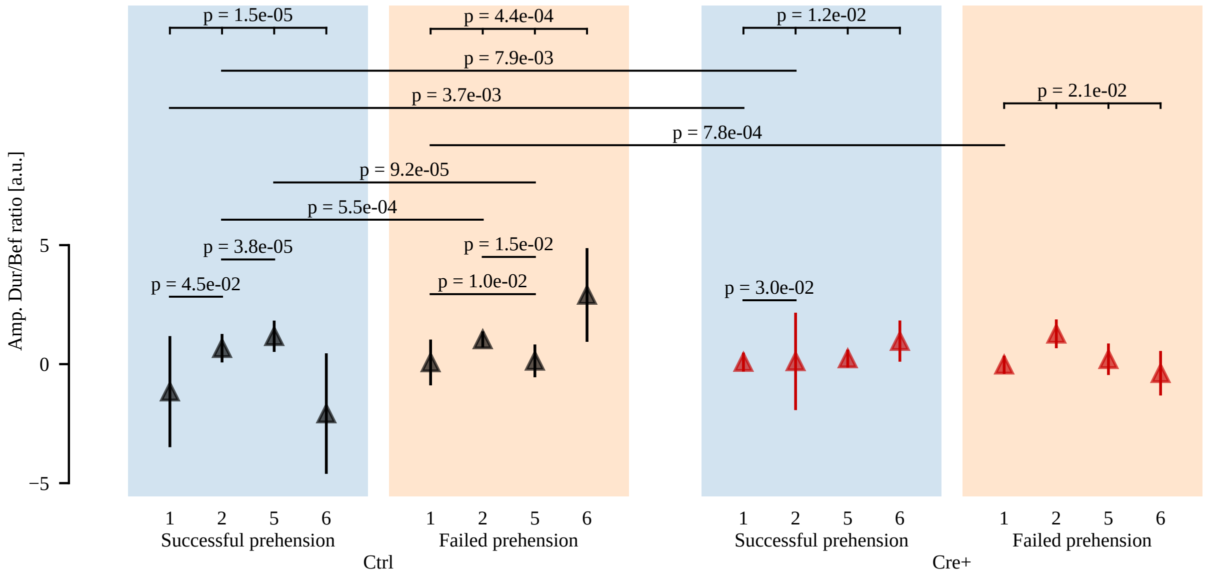


**Supplemental Figure 4:** Planning/execution activity balance is modulated by Ma2 in early sessions. Amplitude ratio of PC activity between the during and before epochs represented as mean ± SEM, separated by genotype (control, Ctrl, black; Cre+, red), prehension accuracy (successful prehension, blue shade; failed prehension, orange shade) and session.

### PC activity peak latency is strongly affected by prehension accuracy

We found a significant isolated effect of prehension accuracy for both prehension phases during all sessions, except for the learning session ($H(1) > 16.552$, $p < 4.7e{-}05$). However, in the learning session, prehension accuracy interacted with genotype ($H(1) > 10.364$, $p < 9.0e{-}03$). These results indicate that PCs strongly modify their temporal activity pattern according to prehension accuracy, while differently affected by genotype, training session, and the phase in which neurons reach their response peak (Supplemental Figure 5).

In agreement with the three-way interaction effect found between prehension accuracy, genotype, and training session on PC activity peak latency (Scheirer-Ray-Hare, $H(3) = 27.514$, $p = 4.6e{-}06$) and the two-way interaction between training session and prehension phase ($H(3) = 20.244$, $p = 1.5e{-}04$), PCs that reached their activity peak before prehensions had decreased latencies in successful prehensions compared to failed prehensions for both genotypes at all sessions (Mann-Whitney U, eff.size $> 0.26$, $p < 3.7e{-}03$; Supplemental Figure 5), except for Chrna2-Cre+ mice in the naive session. Conversely, PCs that reached their activity peak during prehensions had increased latencies in successful prehensions compared to failed prehensions for both genotypes at all sessions (eff.size $> 0.304$, $p < 9.6e{-}04$), except for Chrna2-Cre+ mice in the training session. Since neurons were sorted by latency on successful prehensions at each session, significance bars for comparisons between prehension phases were omitted for successful prehensions.

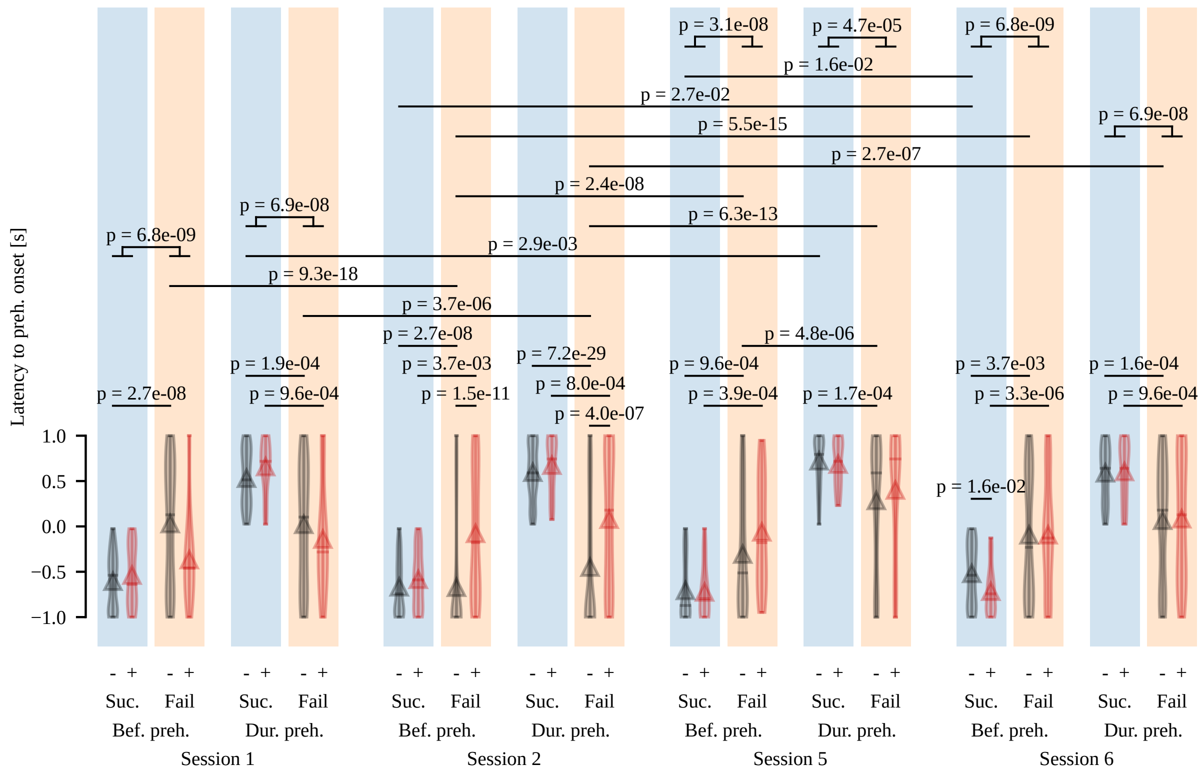


**Supplemental Figure 5:** Distribution of PCs activity peak latencies differ between correct and incorrect prehensions. PC peak latency for successful and failed prehensions, separating neurons by session, the phase of their peak of activity (Bef. preh., before; Dur. preh., during prehension), prehension accuracy (Suc., successful; Fail, failed) and genotype (-, Controls; +, Chrna2-Cre+).

### PC activity sharpness is differently affected by training upon increased M$\alpha$2 cell excitation

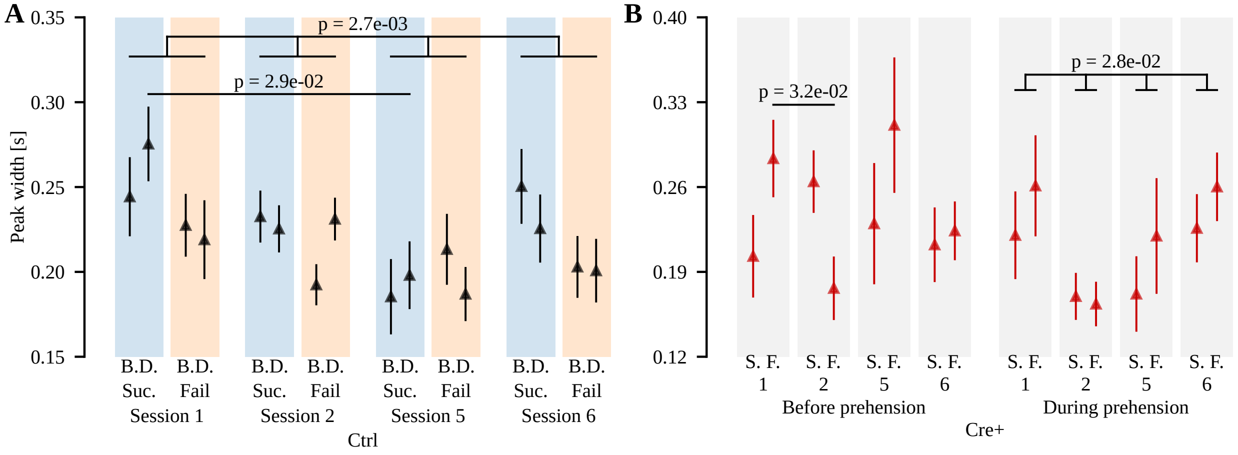


**Supplemental Figure 6:** Training-induced changes in PC activity sharpness are affected by M$\alpha$2 cell increased excitation. A-B) PCs response peak width separated by genotype, session, prehension accuracy (Success, S.; Failed; F.) and prehension phase (Before, B.; During, D.) where the activity peak is reached. The order of factors are different for each genotype in order to highlight the effects of training found in PC peak width.

For control mice, the width of activity peaks from all PCs steadily decreases with training, and slightly increases upon retraining ($H(3) = 15.639$, $p = 2.7e-03$, Supplemental Figure 6A), an effect not observed in Chrna2-Cre+ mice. Although an effect of training session is present for Chrna2-Cre+ mice only for PCs with activity peaks during prehensions ($H(3) = 11.524$, $p = 2.8e-02$, Supplemental Figure 6B), the changes follow a different trend, with peak widths decreasing in the learning session and then increasing on the training and retraining sessions.

### LFP power is overall reduced in non-prehension epochs

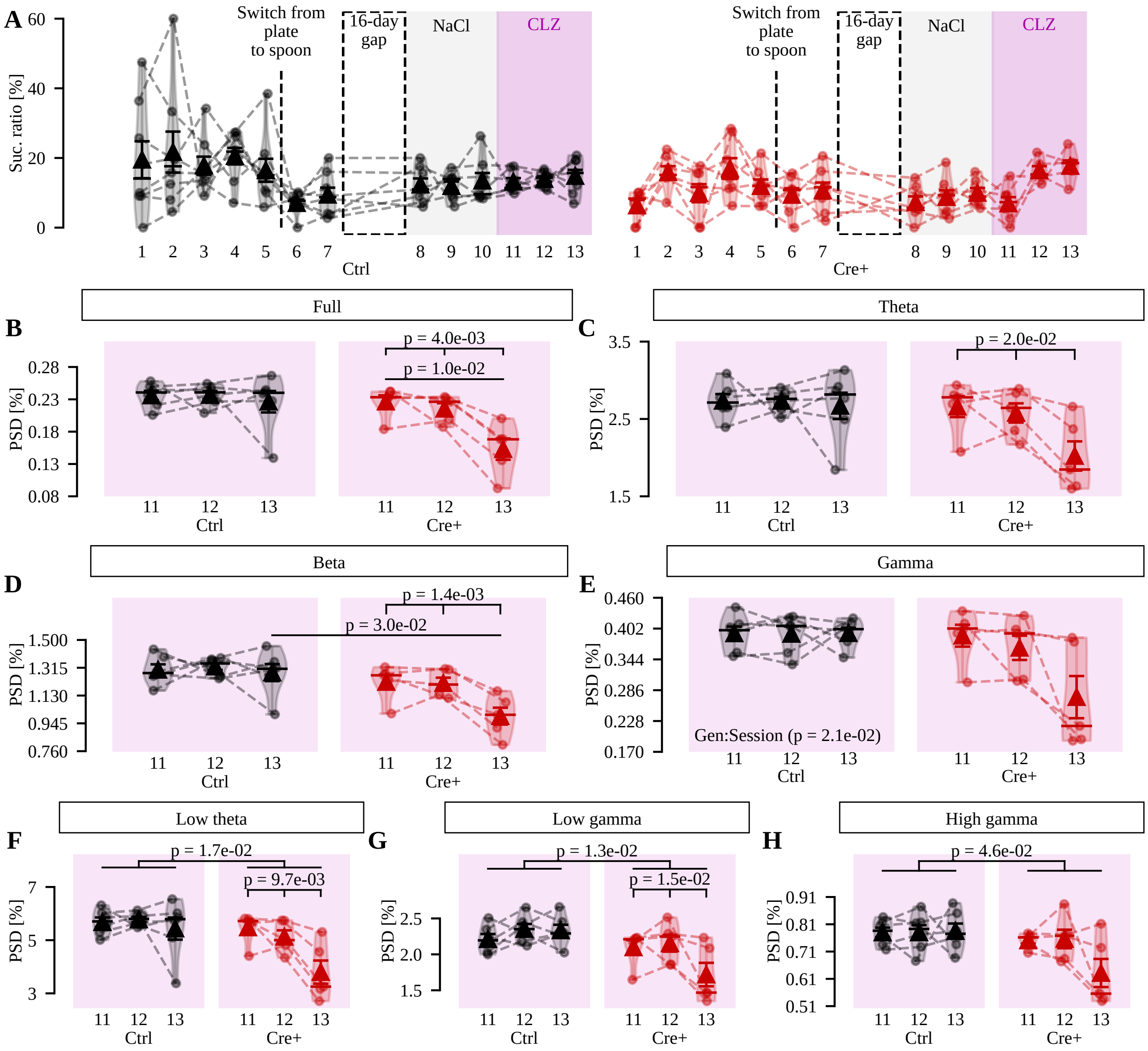


**Supplemental Figure 7:** LFP power is overall reduced in non-prehension epochs. A) Full behavioral sessions indicating the switch from the spoon to the plate pellet delivery and the 16-day gap due to viral injection and implantation procedures. B-E) Relative power spectrum density (PSD) for the full theta, beta and gamma bands, separated by genotype and session. F-H) Same as A-D) but for the subdivisions low theta, low gamma and high gamma.

### Expanded statistical results

**Supplemental Table 1:** Expanded statistical results for behavioral tests. preh.: prehension; Class: prehension accuracy; Gen: genotype; Phase: prehension phase; Session: training session.

| | |
|---|---|
| A | Gen effect on total prehensions: ANOVA, $F(1,11) = 4.8e\text{-}02$, $p = 0.83$ |
| B | Gen effect on successful prehensions: ANOVA, $F(1,11) = 3.4e\text{-}02$, $p = 0.856$ |
| C | Gen effect on success ratio: ANOVA, $F(1,11) = 0.336$, $p = 0.574$ |
| D | Session effect on total prehensions: ANOVA, $F(10,110) = 9.525$, $p = 2.9e\text{-}11$ |
| E | Session effect on successful prehensions: ANOVA, $F(10,110) = 7.571$, $p = 4.3e\text{-}09$ |
| F | Session effect on success ratio: ANOVA, $F(10,110) = 5.536$, $p = 1.2e\text{-}06$ |
| G | Overall session effect: ANOVA, $F(6,54) = 4.27$, $p = 1.0e\text{-}03$ |
| H | Overall Gen:Session interaction effect: ANOVA, $F(6,54) = 2.203$, $p = 0.057$ |
| I | Session effect for Chrna2-Cre+ mice: ANOVA, $F(6,24) = 5.612$, $p = 1.9e\text{-}03$ |
| J | Session effect for Control mice: ANOVA, $F(6,30) = 1.083$, $p = 0.395$ |
| K | Treatment effect for Chrna2-Cre+ mice: ANOVA, $F(1,15) = 10.974$, $p = 1.0e\text{-}02$ |
| L | Treatment effect for Control mice: ANOVA, $F(1,19) = 0.878$, $p = 0.361$ |

**Supplemental Table 2:** Expanded statistical results for neuronal activity amplitude. act.: activity; preh.: prehension; Class: prehension accuracy; Gen: genotype; Phase: prehension phase; Session: training session.

| | |
|---|---|
| A | Class:Session effect in PC act. amplitude; Scheirer-Ray-Hare, $H(3) = 10.378$, $p = 1.6e\text{-}02$ |
| B | Gen:Session effect in PC act. amplitude during preh.; Scheirer-Ray-Hare, $H(3) = 12.057$, $p = 7.2e\text{-}03$ |
| C | Kruskal-Wallis, $F(3,844) = 4.951$, $p = 8.2e\text{-}03$ |
| D | Mann-Whitney U eff. size = 0.147; $p = 5.5e\text{-}03$ |
| E | Mann-Whitney U eff. size = 0.154; $p = 1.0e\text{-}02$ |
| F | Scheirer-Ray-Hare, $H(1) = 10.497$, $p = 4.8e\text{-}03$ |
| G | ANOVA-type, F2-LD-F1 model $F(1,inf) = 8.623$, $p = 1.3e\text{-}02$ |
| H | Kruskal-Wallis, $F(1,535) = 9.446$, $p = 3.6e\text{-}02$ |
| I | Gen effect in PC act. amplitude; ANOVA-type, F2-LD-F1 model $F(1,inf) > 5.846$, $p < 4.7e\text{-}02$ |
| J | Class:Session effect in PC act. amplitude of controls; Scheirer-Ray-Hare, $H(3) = 13.654$, $p = 6.8e\text{-}03$ |
| K | Scheirer-Ray-Hare, $H(3) = 23.752$, $p = 2.8e\text{-}05$ |
| L | Gen effect for successful prehensions on session 1; Kruskal-Wallis, $F(1,212) = 12.418$, $p = 3.6e\text{-}03$ |
| M | Gen effect for failed prehensions on session 1; Kruskal-Wallis, $F(1,266) = 15.674$, $p = 7.7e\text{-}04$ |
| N | Gen effect for successful prehensions on session 2; Kruskal-Wallis, $F(1,479) = 10.449$, $p = 7.9e\text{-}03$ |
| O | Class effect for Controls, session 2; Kruskal-Wallis, $F(1,535) = 15.857$, $p = 5.4e\text{-}04$ |
| P | Class effect for Controls, session 5; Kruskal-Wallis, $F(1,574) = 19.586$, $p = 9.2e\text{-}05$ |

**Supplemental Table 3:** Expanded statistical results for neuronal activity latency. act.: activity; preh.: prehension; Class: prehension accuracy; Gen: genotype; Phase: prehension phase; Session: training session.

| | |
|---|---|
| A | Class:Gen:Session effect in PC act. peak latency; Scheirer-Ray-Hare, $H(3) = 27.514$, $p = 4.6e\text{-}06$ |
| B | Phase:Session effect in PC act. peak latency; Scheirer-Ray-Hare, $H(3) = 20.244$, $p = 1.5e\text{-}04$ |
| C | Class effect in PC act. peak latency for both prehension phases at all sessions except session 2; Scheirer-Ray-Hare, $H(1) > 16.552$, $p < 4.7e\text{-}05$ |
| D | Mann-Whitney U eff. size = 0.371, $p = 4.0e\text{-}07$ |
| E | Mann-Whitney U eff. size = 0.277, $p = 1.6e\text{-}02$ |

**Supplemental Table 4:** Expanded statistical results for neuronal activity peak width. act.: activity; preh.: prehension; Class: prehension accuracy; Gen: genotype; Phase: prehension phase; Session: training session.

| | |
|---|---|
| A | Class:Gen:Session effect in PC act. peak width; Scheirer-Ray-Hare, $H(3) = 10.028$, $p = 1.8e-02$ |
| B | Class:Phase:Session effect in PC act. peak width; Scheirer-Ray-Hare, $H(3) = 10.612$, $p = 1.4e-02$ |
| C | Gen:Phase:Session effect in PC act. peak width; Scheirer-Ray-Hare, $H(3) = 17.698$, $p = 5.1e-04$ |
| D | Gen effect in PC act. peak width during successful preh.; Scheirer-Ray-Hare, $H(1) = 8.86$, $p = 1.2e-02$ |
| E | Gen effect in PC act. peak width during session 2; Scheirer-Ray-Hare, $H(1) = 16.629$, $p = 3.6e-04$ |

**Supplemental Table 5:** Expanded statistical results for neuronal assemblies. act.: activity; preh.: prehension; Class: prehension accuracy; Gen: genotype; Phase: prehension phase; Session: training session.

| | |
|---|---|
| A | ANOVA-type, F2-LD-F1 model $F(2,inf) = 5.50$, $p = 4.7e-03$ |
| B | Mann-Whitney U eff. size = 0.55, $p = 3.6e-03$ |
| C | Mann-Whitney U eff. size = 0.42, $p = 4.5e-02$ |
| D | Gen effect in assembly area; Scheirer-Ray-Hare, $H(1) = 10.539$, $p = 1.2e-03$ |
| E | Mann-Whitney U eff. size = 0.495, $p = 1.2e-02$ |
| F | Gen:Session effect in assembly resilience; ANOVA, $F(2,94) = 4.18$, $p = 1.8e-02$ |
| G | t-test eff. size = 0.937, $p = 1.2e-02$ |

**Supplemental Table 6:** Expanded statistical results for LFP recordings. preh.: prehension; Class: prehension accuracy; Gen: genotype; Phase: prehension phase; Session: training session.

| | |
|---|---|
| A | Gen effect in theta power; ANOVA, $F(1,8) = 6.783$, $p = 3.1e-02$ |
| B | Class effect in theta power; ANOVA, $F(1,8) = 6.332$, $p = 3.6e-02$ |
| C | Gen effect effect in low theta power; ANOVA, $F(1,8) = 6.489$, $p = 3.4e-02$ |
| D | Class effect effect in high theta power; ANOVA, $F(1,8) = 8.724$, $p = 1.8e-02$ |
| E | Class:Gen:Phase effect in gamma power; ANOVA-type, F1-LD-F2 model $F(1,inf) = 5.919$, $p = 4.5e-02$ |
| F | Class effect in session 13 gamma power for Chrna2-Cre+ before preh.; ANOVA-type, LD-F1 model $F(1,inf) = 8.333$, $p = 4.3e-02$ |
| G | Gen effect in session 13 low gamma power; ANOVA-type, F1-LD-F2 model $F(1,inf) = 8.781$, $p = 9.1e-03$ |
| H | Class effect in session 13 low gamma power for Chrna2-Cre+ before preh.; ANOVA-type, LD-F1 model $F(1,inf) = 8.333$, $p = 4.3e-02$ |
| I | Gen:Session effect in session 13 high gamma power; ANOVA, $F(2,16) = 3.706$, $p = 4.8e-02$ |
| J | Gen effect in session 13 high gamma power; ANOVA, $F(1,8) = 9.329$, $p = 4.8e-02$ |
| K | ANOVA, $F(2,18) > 3.584$, $p < 4.9e-02$ |
| L | ANOVA, $F(2,8) > 8.667$, $p < 2.0e-02$ |
| M | Gen effect, ANOVA-type, F1-LD-F1 model $F(1,inf) = 5.687$, $p = 1.7e-02$ |
| N | Session effect, ANOVA-type, F1-LD-F1 model $F(2,inf) = 3.37$, $p = 3.7e-02$ |
| O | Low-gamma, ANOVA-type, F1-LD-F1 model $F(1,inf) = 6.225$, $p = 1.3e-02$ |
| P | High-gamma, ANOVA, $F(1,9) = 5.356$, $p = 4.6e-02$ |
